# Characterization and validation of 16 α-synuclein conformation-specific antibodies using well-characterized preparations of α-synuclein monomers, fibrils and oligomers with distinct structures and morphology: How specific are the conformation-specific α-synuclein antibodies?

**DOI:** 10.1101/2020.06.15.151514

**Authors:** Senthil T. Kumar, Somanath Jagannath, Cindy Francois, Hugo Vanderstichele, Erik Stoops, Hilal A. Lashuel

**Author notes:** To whom correspondence should be addressed: Laboratory of Molecular and Chemical Biology of Neurodegeneration, Brain Mind Institute, Ecole Polytechnique Fédérale de Lausanne, 1015 Lausanne. These authors contributed equally to the manuscript (Senthil T. Kumar, Somanath Jagannath).

## Abstract

Increasing evidence suggests that alpha-synuclein (α-syn) oligomers are obligate intermediates in the pathway involved in α-syn fibrillization and Lewy body (LB) formation, and may also accumulate within LBs in Parkinson’s disease (PD) and other synucleinopathies. Therefore, the development of tools and methods to detect and quantify α-syn oligomers has become increasingly crucial for mechanistic studies to understand the role of these oligomers in PD, and to develop new diagnostic methods and therapies for PD and other synucleinopathies. The majority of these tools and methods rely primarily on the use of aggregation state-specific or conformation-specific antibodies. Given the impact of the data and knowledge generated using these antibodies on shaping the foundation and directions of α-syn and PD research, it is crucial that these antibodies are thoroughly characterized, and their specificity or ability to capture diverse α-syn species is tested and validated. Herein, we describe an antibody characterization and validation pipeline that allows a systematic investigation of the specificity of α-syn antibodies using well-defined and well-characterized preparations of various α-syn species, including monomers, fibrils, and different oligomer preparations that are characterized by distinct morphological, chemical and secondary structure properties. This pipeline was used to characterize 18 α-syn antibodies, 16 of which have been reported as conformation- or oligomer-specific antibodies, using an array of techniques, including immunoblot analysis (slot blot and Western blot), a digital ELISA assay using single molecule array technology and surface plasmon resonance. Our results show that i) none of the antibodies tested are specific for one particular type of α-syn species, including monomers, oligomers or fibrils; ii) all antibodies that were reported to be oligomer-specific also recognized fibrillar α-syn; and iii) a few antibodies showed high specificity for oligomers and fibrils but did not bind to monomers. These findings suggest that the great majority of α-syn aggregate-specific antibodies do not differentiate between oligomers and fibrils, thus highlighting the importance of exercising caution when interpreting results obtained using these antibodies. Our results also underscore the critical importance of the characterization and validation of antibodies before their use in mechanistic studies and as diagnostic and therapeutic agents. This will not only improve the quality and reproducibility of research and reduce costs but will also reduce the number of therapeutic antibody failures in the clinic.

## Introduction

Several neurodegenerative disorders are characterized by the presence of cytoplasmic proteinaceous inclusions termed Lewy bodies (LBs), which are enriched in misfolded and aggregated forms of the presynaptic protein alpha-synuclein (α-syn) (Goedert *et al*. 2017). These diseases include Parkinson’s disease (PD), dementia with Lewy bodies (DLB), and multiple system atrophy (MSA), which are collectively referred to as synucleinopathies. Early studies of the ultrastructural properties and compositions of LBs revealed that they are highly enriched in filamentous structures (Duffy & Tennyson 1965; Lashuel 2020), which were later shown to be composed of α-syn (Spillantini *et al*. 1997; Spillantini *et al*. 1998). These findings, combined with the discovery that mutations in the gene that encodes α-syn causes early-onset forms of PD (Polymeropoulos *et al*. 1997), led to the hypothesis that the process of α-syn fibrillisation and LB formation plays a central role in the pathogenesis of PD and other synucleinopathies. However, the failure of this hypothesis to explain several neuropathological and experimental observations prompted the possibility that intermediates generated on the pathway to α-syn fibrillization and LB formation, rather than the fibrils or LBs themselves, are the primary toxicity-inducing and disease-causing species. These observations include 1) the lack of a strong correlation between Lewy pathology burden, neurodegeneration and disease severity (Colosimo *et al*. 2003; Parkkinen *et al*. 2008); 2) the presence of LBs in the brains of individuals who do not show any symptoms of PD or other synucleinopathies at the time of death (Parkkinen *et al*. 2005; Frigerio *et al*. 2011); and 3) the identification of patients who exhibit Parkinsonian symptoms in the absence of LBs e.g. PD patients harboring parkin and LRRK2 G2019S mutations (Kay *et al*. 2005; Gaig *et al*. 2006; Cookson *et al*. 2008; Johansen *et al*. 2018). These observations are similar to those demonstrating the lack of a correlation between amyloid-plaque burden and cognitive decline in Alzheimer’s disease (AD) (Nelson *et al*. 2012; Jung *et al*. 2016; Arboleda-Velasquez *et al*. 2019), which have supported the toxic oligomer hypothesis of AD.

Several lines of evidence support the α-syn oligomer hypothesis. Both on- and off-pathway soluble and nonfibrillar α-syn oligomers of different sizes and morphologies were consistently observed during the *in vitro* aggregation of α-syn under different conditions (Conway *et al*. 2000; Lashuel *et al*. 2002; Cappai *et al*. 2005). Subsequent studies over the past decade have also provided evidence for the presence of α-syn oligomers in biological fluids such as saliva, blood plasma, basal tears, and cerebrospinal fluid (CSF) from patients suffering from PD and other synucleinopathies (El-Agnaf *et al*. 2006; Tokuda *et al*. 2010; Hirohata *et al*. 2011; Wang *et al*. 2011; Majbour *et al*. 2016; Vivacqua *et al*. 2016; Hamm-Alvarez *et al*. 2019). Several of these studies suggested that the level of oligomers is correlated with the diagnosis of PD and/or disease progression (Sharon *et al*. 2003; El-Agnaf *et al*. 2006; Paleologou *et al*. 2009). One major caveat of these studies is that they were potentially carried out using tools and immunoassays that do not distinguish between oligomers and other higher-order aggregated forms of α-syn (fibrils or amorphous aggregates). Nonetheless, they paved the way for further studies demonstrating that α-syn oligomers/aggregates are secreted by neurons (Sharon *et al*. 2003; Tofaris *et al*. 2003; Jang *et al*. 2010; Tokuda *et al*. 2010; Majbour *et al*. 2016) in the brain, and could mediate the propagation of α-syn pathology and cause neurodegeneration. Indeed, several studies have reported that α-syn oligomers are released by neurons via exocytosis (Jang *et al*. 2010) and are then taken up by other cells via different mechanisms, including endocytosis (Desplats *et al*. 2009), trans-synaptic propagation (Danzer *et al*. 2012) or receptor-mediated uptake (Lee *et al*. 2008). Furthermore, α-syn oligomers have been shown to directly or indirectly contribute to α-syn-induced toxicity and neurodegeneration via different mechanisms, including but not limited to i) the disruption of cell membrane integrity by the formation of pores in the membrane (Volles *et al*. 2001; Danzer *et al*. 2007); ii) synaptic toxicity or neuronal signaling dysfunction (Diógenes *et al*. 2012; Rockenstein *et al*. 2014; Kaufmann *et al*. 2016; van Diggelen *et al*. 2019); iii) the failure of protein degradation pathways (Cuervo *et al*. 2004; Klucken *et al*. 2012; Tekirdag & Cuervo 2018); iv) endoplasmic reticulum dysfunction (Colla *et al*. 2012); v) mitochondrial dysfunction (Parihar *et al*. 2009; Di Maio *et al*. 2016); and vi) the enhancement of inflammatory responses (Wilms *et al*. 2009). These observations, combined with the overwhelming evidence that oligomer-induced toxicity is a key contributor or driving force leading to neurodegeneration in Alzheimer’s disease (AD), fueled greater interest in the development of tools, therapies and diagnostics that specifically target α-syn oligomers. This includes the development of various protocols for the preparation of oligomers, the generation of oligomer-specific antibodies, and immunoassays for quantifying oligomers.

Oligomers can be broadly defined as all the soluble multimeric species that exist before the formation of α-syn fibrils, including a) dimers, trimers and low molecular weight assemblies, which are not easily discernable by electron microscopy (EM) and atomic force microscopy (AFM)), and b) higher molecular weight oligomers with different morphologies that are composed of >10 monomers, which are easily detectable by EM, AFM and other imaging techniques (Lashuel *et al*. 2002; Lashuel & Lansbury 2006; Stöckl *et al*. 2013; Cremades *et al*. 2017; Ruggeri *et al*. 2018; Kumar *et al*. 2020a). Our current knowledge of the biophysical properties of α-syn oligomers has been shaped primarily by results obtained by investigating α-syn aggregation and fibril formation *in vitro*. The propensity of α-syn to form oligomers is highly dependent on several factors, such as the protein concentration and sequence (including the presence of disease-associated mutations and post-translational modifications) (Lashuel *et al*. 2002; Paslawski *et al*. 2014a; Paslawski *et al*. 2014b), interactions with metals, other proteins and small molecules, and chemical modification by specific molecules (e.g. dopamine, 4-oxo-2-nonenal, 4-hydroxy-2-nonenal (HNE), and epigallocatechin gallate) (Danzer *et al*. 2007; Qin *et al*. 2007; Ehrnhoefer *et al*. 2008; Näsström *et al*. 2011b). Depending on the conditions used, different types of α-syn oligomers have been consistently observed *in vitro*, including globular, spherical, amorphous, curvilinear and pore-like oligomers (Figure 1) (Lashuel *et al*. 2002; Kumar *et al*. 2020a). It remains unknown to what extent these oligomers resemble the oligomers that form in different cell types in the brains of patients. Several studies have reported the detection of oligomers in cell cultures, in the brains of animal models of synucleinopathies, and during the analysis of cerebrospinal fluids (CSF) and postmortem examinations of brains of PD, DLB and MSA patients (Sharon *et al*. 2003; Tofaris *et al*. 2003; Jang *et al*. 2010; Tokuda *et al*. 2010; Majbour *et al*. 2016). However, the evidence to support the presence of specific oligomers in these studies has been based for the most part on the detection of SDS-resistant oligomeric bands by Western blotting (Baba *et al*. 1998; Sharon *et al*. 2003; Tsigelny *et al*. 2008), the use of proximity ligation assays (Roberts *et al*. 2015), or the reliance on “oligomer-specific” antibodies or immunoassays. Thus, much of the knowledge and many of the hypotheses in the field today are based on conclusions drawn from studies relying on antibodies.

**Figure 1:**
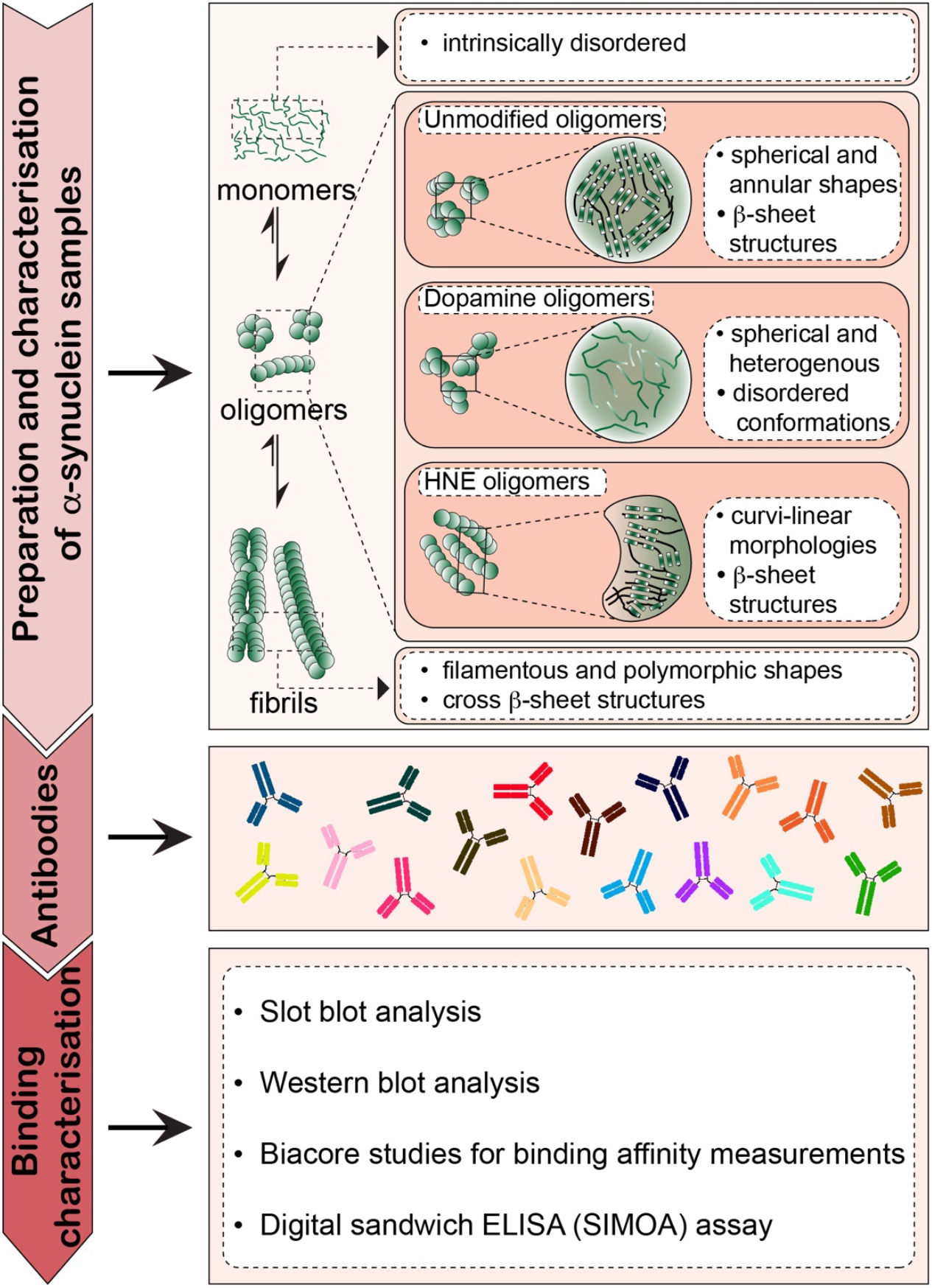
A schematic illustration of our antibody validation strategy. In brief, the pipeline included the preparation of well-defined preparations of α-syn monomers, oligomers, and fibrils. Oligomers were generated from three different protocols in an attempt to partially capture the morphological, chemical and structural heterogeneity of oligomers *in vivo*. The α-syn conformation-specific antibodies were procured from different sources. The immunoreactivity of these antibodies was assessed using slot blot, Western blot, a digital sandwich ELISA (SIMOA) assays and SPR.

One major untested assumption related to the use of oligomer-specific antibodies and immunoassays is that the antibodies used are capable of capturing the structural and morphological diversity of α-syn oligomers *in vivo*. Notably, all these antibodies were generated using specific recombinant α-syn aggregates, fibrils or oligomers generated under *in vitro* conditions. Some of the limitations of existing antibody validation approaches include the following: 1) the lack of detailed characterization of *in vitro* oligomer preparations with respect to their purity, homogeneity and structural properties; 2) the use of oligomer preparations that may not reflect the conformational, biochemical and morphological diversity that exists in the brain; and 3) the lack of research that establishes whether the specificity of antibodies is driven by their high affinity for oligomers, or by the avidity binding characteristics of the antibodies.

Given the impact of the use of antibodies on shaping our knowledge of α-syn and its role in health and disease, and on developing diagnostics and therapies for PD and synucleinopathies, we developed a protocol that enables systematic assessment of the specificity of α-syn antibodies using well-defined and well-characterized preparations of α-syn fibrils, oligomers, and monomers. This approach was then used to evaluate a library of 18 α-syn antibodies, 16 of which were reported to be aggregate-specific (Table 2). These antibodies can be broadly classified depending on the immunogens used for their generation: oligomers based on the use of i) a modified version of fulllength α-synuclein (antibody clones 24H6, 12C6, 26F1, and 26B10); ii) *in vitro* generated α-syn fibrils (antibody clones 7015, 9029, SYNO2, SYNO3, and SYNO4); iii) recombinant α-syn aggregates (antibody clones A17183A, A171183B, A17183E, and A17183G); iv) synthetic α-syn peptides encompassing amino acids 44 to 57 (5G4) or filaments derived from recombinant “exact sequence is not disclosed by the vendor” (MJFR-14) or amino acids 115-125 (ASyO5); vi) recombinant full-length α-syn monomers (SYN211); and vi) a recombinant truncated α-syn variant consisting of residues 15-123 (SYN-1) (Table 2). To verify the specificity of these antibodies, we first screened them all against well-characterized preparations of α-syn species (monomers, oligomers, and fibrils) using immunoblot analysis (slot blotting and Western blotting) and a digital enzyme-linked immunosorbent assay (ELISA) using single molecule array (SIMOA) technology (Figure 1). To further scrutinize the conformational specificity of the antibodies, we tested them against different types of oligomer that exhibit distinct morphological, chemical and secondary structure properties. Finally, the binding affinity of selected antibodies was determined using surface plasmon resonance (SPR). This approach enabled us to define the specificity of the antibodies to a high degree and show that although some antibodies were specific for aggregated forms of α-syn and did not recognize monomers, all antibodies that were reported to be oligomer-specific also recognized fibrillar α-syn. Furthermore, some of the antibodies that were reported to be oligomer- or fibril-specific also recognized α-syn monomers. We also identified an antibody that showed a preference for β-sheet-enriched fibrils and oligomers, but not for disordered oligomers or monomers. Our studies reveal that none of the antibodies tested (Table 2) showed any unique preferential specificity for one particular form of α-syn species, including monomers, oligomers or fibrils, and that it is possible to develop antibodies that recognize diverse α-syn oligomers and fibrils. Our work underscores the importance of using well-characterized tools (*in vitro*-produced calibrants) and multiple methods to define the specificity of antibodies. This will not only help us to advance PD research, but will also improve the selection of promising antibody candidates and reduce the number of failures in advanced clinical trials of PD therapeutics.

**Table 1:**
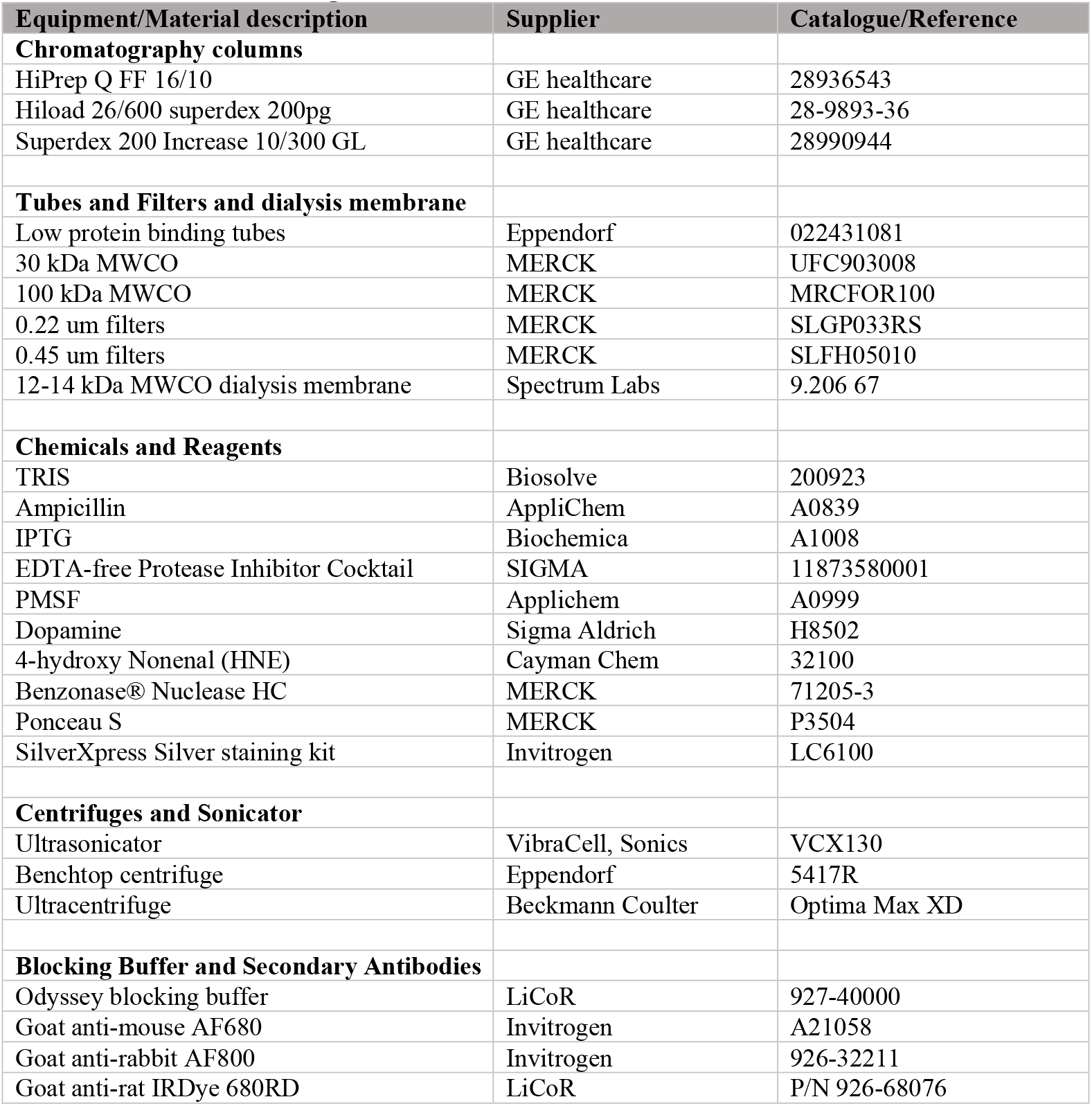
Materials and Reagents

| **Equipment/Material description** | **Supplier** | **Catalogue/Reference** |
| --- | --- | --- |
| **Chromatography columns** |  |  |
| HiPrep Q FF 16/10 | GE healthcare | 28936543 |
| Hiload 26/600 superdex 200pg | GE healthcare | 28-9893-36 |
| Superdex 200 Increase 10/300 GL | GE healthcare | 28990944 |
| **Tubes and Filters and dialysis membrane** |  |  |
| Low protein binding tubes | Eppendorf | 022431081 |
| 30 kDa MWCO | MERCK | UFC903008 |
| 100 kDa MWCO | MERCK | MRCFOR100 |
| 0.22 um filters | MERCK | SLGP033RS |
| 0.45 um filters | MERCK | SLFH05010 |
| 12-14 kDa MWCO dialysis membrane | Spectrum Labs | 9.206 67 |
| **Chemicals and Reagents** |  |  |
| TRIS | Biosolve | 200923 |
| Ampicillin | AppliChem | A0839 |
| IPTG | Biochemica | A1008 |
| EDTA-free Protease Inhibitor Cocktail | SIGMA | 11873580001 |
| PMSF | Applichem | A0999 |
| Dopamine | Sigma Aldrich | H8502 |
| 4-hydroxy Nonenal (HNE) | Cayman Chem | 32100 |
| Benzonase® Nuclease HC | MERCK | 71205-3 |
| Ponceau S | MERCK | P3504 |
| SilverXpress Silver staining kit | Invitrogen | LC6100 |
| **Centrifuges and Sonicator** |  |  |
| Ultrasonicator | VibraCell, Sonics | VCX130 |
| Benchtop centrifuge | Eppendorf | 5417R |
| Ultracentrifuge | Beckmann Coulter | Optima Max XD |
| **Blocking Buffer and Secondary Antibodies** |  |  |
| Odyssey blocking buffer | LiCoR | 927-40000 |
| Goat anti-mouse AF680 | Invitrogen | A21058 |
| Goat anti-rabbit AF800 | Invitrogen | 926-32211 |
| Goat anti-rat IRDye 680RD | LiCoR | P/N 926-68076 |

**Table 2:**
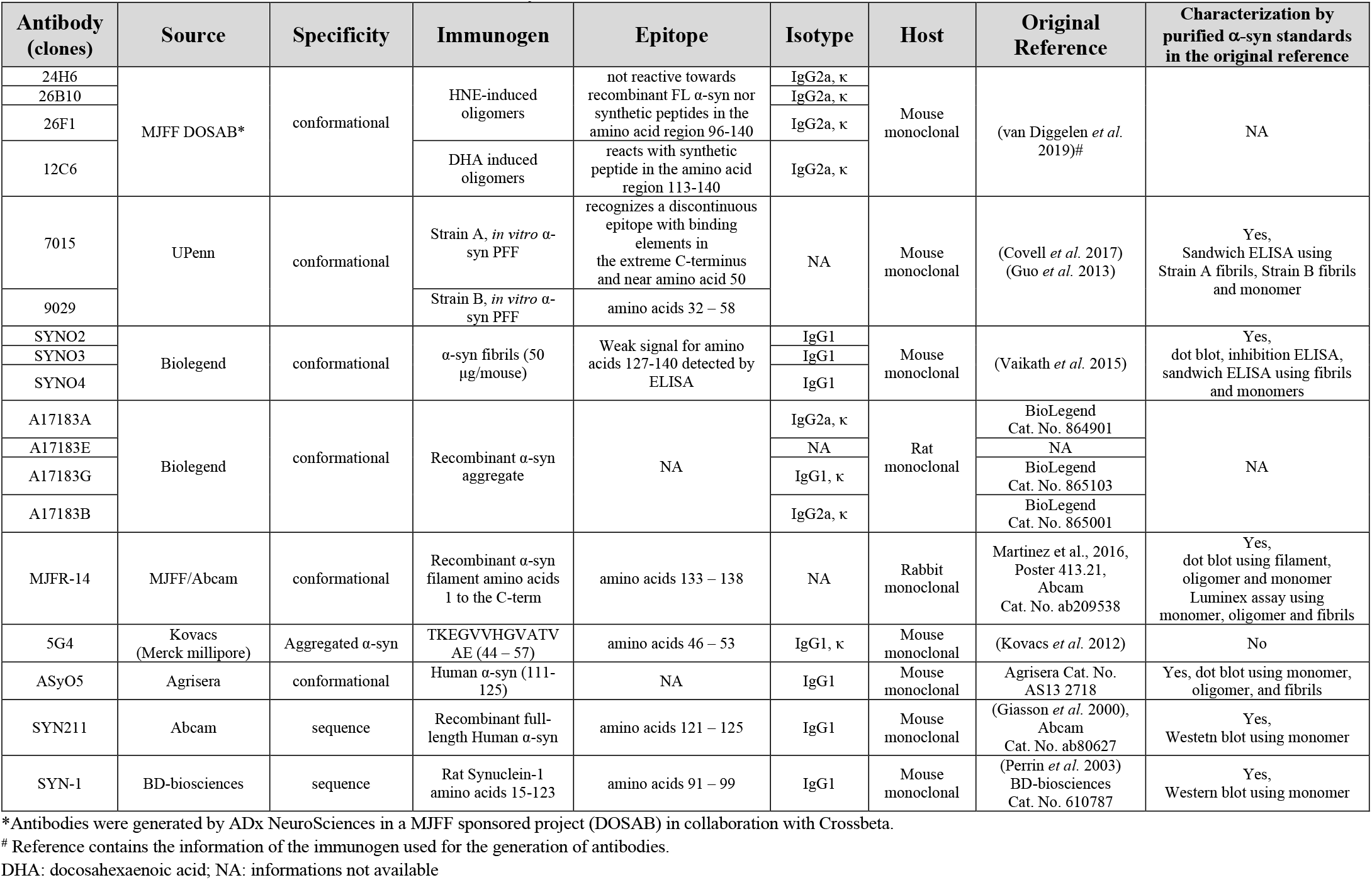
List and details of antibodies used in this study

## Results

### Preparation and characterization of α-syn monomers, oligomers and fibrils

To investigate the specificity of the antibodies listed in Table 2, we first assessed their specificity towards α-syn monomers, oligomers, and fibrils. To accomplish this goal, we generated well-characterized preparations of human α-syn 1) fibrils, 2) oligomers and 3) monomers that were “free” of cross-species contamination. The purity of each preparation was verified using our recently described centrifugation-filtration protocol (Kumar *et al*. 2020a) (Figure 2A). Given that α-syn oligomers and fibrils are always in equilibrium with monomers, it is difficult to eliminate the presence of monomers completely. To eliminate or minimize the amount of monomers (<5%), all fibril and oligomeric samples were subjected to centrifugation-filtration protocol immediately prior to their use in our studies, as previously described (Kumar *et al*. 2020a). Similarly, to ensure that the α-syn monomeric preparations were free of any preformed aggregates, the monomeric samples were filtered through a 100 kDa filter, and the flow-through (aggregate-free monomers) was collected and kept on ice and used immediately.

**Figure 2:**
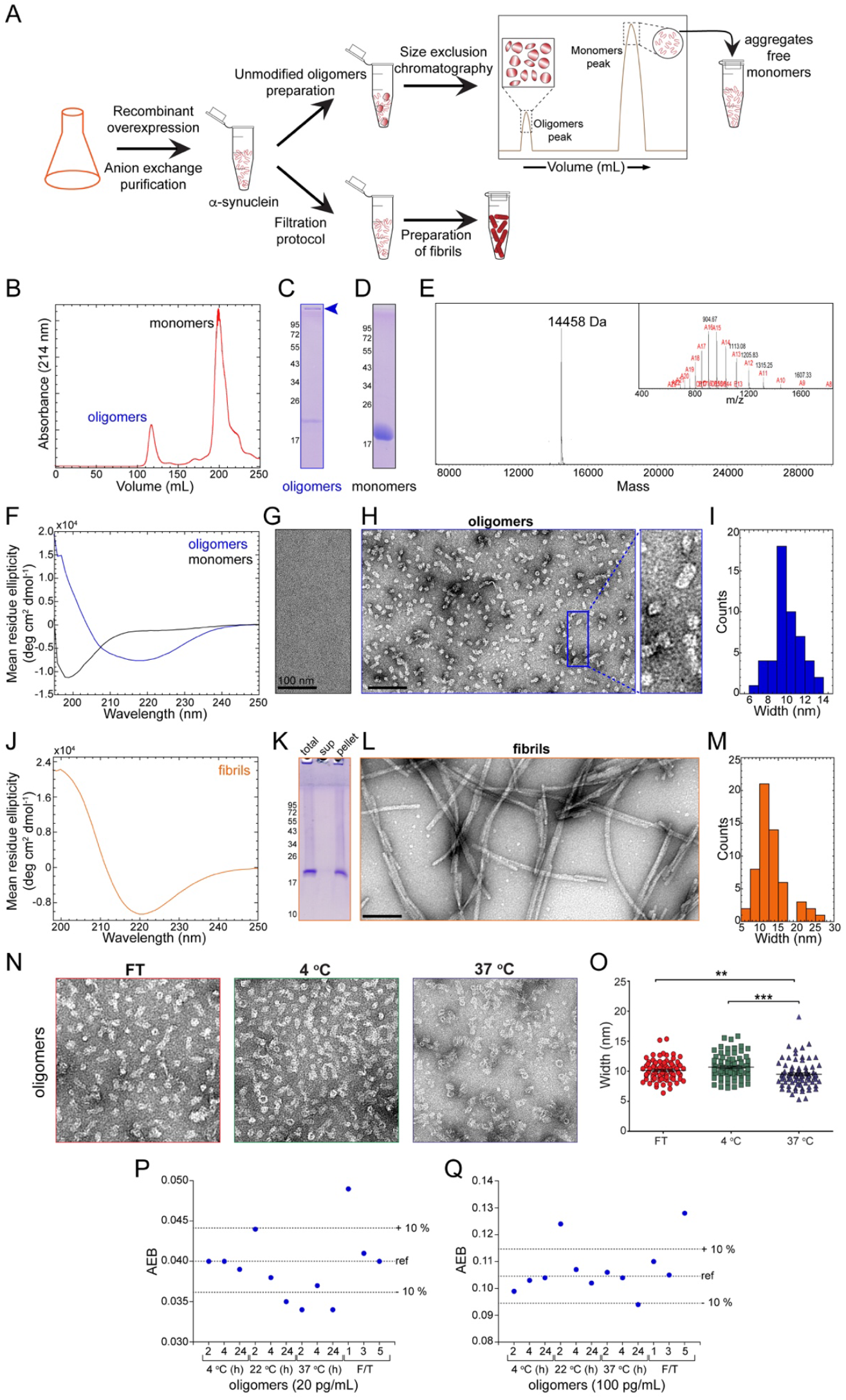
Preparation and characterization of α-syn monomers, oligomers, and fibrils. (A) A scheme depicting the preparation of α-syn monomers, oligomers, and fibrils. (B) SEC purification of monomers and oligomers. (C and D) SDS-PAGE analysis followed by Coomassie staining of oligomers (C) and monomers (D) purified from B. (E) ESI-MS spectra of monomers separated by SEC. (F) Comparison of the CD spectra of monomers and oligomers. Oligomers were predominantly enriched in β-sheet structures and monomers showed a predominantly disordered structure. (G) Negatively stained EM analysis performed on monomers. (H) Negatively stained EM analysis performed on oligomers. (I) Width distribution of oligomers. (J) CD spectra of fibrils. (K) SDS-PAGE analysis followed by Coomassie staining of the total, supernatant, and pellet fractions obtained during fibril preparation. The pellet fraction is the fibril fraction devoid of monomers and oligomers that was used for further binding studies. (L) Negatively stained EM analysis performed on fibrils. (M) Width distribution of the fibrils. (N) EM images and width distribution (O) of oligomers subjected to different temperature conditions, freeze-thawing cycles and incubation at 4 °C and 37 °C. and Q) Assessment of the stability of the oligomeric preparation by ELISA. Two concentrations of oligomers (20 pg/mL (P) and 100 pg/mL (Q)) were incubated at three different temperatures ranges (2-8°C, 20-25°C and 36-38°C) for varying times (2 h, 4 h and 24 h) or subjected to multiple freeze-thaw (F/T) cycles (1x, 3x, and 5x) (X-axis). The signals (AEB) from the immunoassay with the antibody A17183B were plotted on the Y-axis. The dashed lines represent the reference condition “ref” and a 10% decrease/increase in the AEB signal as arbitrary thresholds.

Several procedures have been developed with the aim of generating homogenous preparations of oligomers *in vitro*, but all were shown to result in preparations that contain mixtures of oligomers that are structurally and morphologically diverse. However, it is possible to generate preparations that are enriched in specific oligomeric species by subfractionating these preparations using size exclusion chromatography or other separation methods (Lashuel *et al*. 2002; Lashuel & Lansbury 2006). The protocols involve the generation of oligomers either by incubating recombinant α-syn monomers at high concentrations in buffers with or without additional components such as dopamine (Conway *et al*. 2001; Cappai *et al*. 2005; Norris *et al*. 2005; Leong *et al*. 2009; Rekas *et al*. 2010; Choi *et al*. 2013; Planchard *et al*. 2014), lipids (Broersen *et al*. 2006; Trostchansky *et al*. 2006; Qin *et al*. 2007; Nasstrom *et al*. 2009; Näsström *et al*. 2011a; Näsström *et al*. 2011b; De Franceschi *et al*. 2011; Diógenes *et al*. 2012; Xiang *et al*. 2013), metals (Lowe *et al*. 2004; Cole *et al*. 2005; Danzer *et al*. 2007; Danzer *et al*. 2009; Wright *et al*. 2009; Schmidt *et al*. 2012), alcohols (Danzer *et al*. 2007; Ehrnhoefer *et al*. 2008; Danzer *et al*. 2009; Illes-Toth *et al*. 2015), or by using methods that are based on the use of chemical cross-linking agents (Ruesink *et al*. 2019). In the absence of additional components, oligomers are found to exhibit heterogeneous morphologies, such as globular, spherical, annular pore-shaped, rectangular and tubular-shaped, and are usually but not always enriched in β-sheet structures (Lashuel *et al*. 2002). In the presence of additional components such as dopamine, lipids or alcohols, oligomers are found to have spherical, globular, rod-shaped or curvilinear morphologies, which are structurally different from primarily disordered, α-helical or β-sheeted structures, suggesting that the formation of oligomers is strongly influenced by the environment in which they form (Conway *et al*. 2001; Lowe *et al*. 2004; Norris *et al*. 2005; Broersen *et al*. 2006; Danzer *et al*. 2007; Nasstrom *et al*. 2009; Rekas *et al*. 2010; De Franceschi *et al*. 2011; Näsström *et al*. 2011a; Diógenes *et al*. 2012; Bae *et al*. 2013; Choi *et al*. 2013; Fecchio *et al*. 2013; Planchard *et al*. 2014).

Recombinant α-syn was used for the preparation of oligomers and fibrils. For the preparation of oligomers (Lashuel *et al*. 2002; Paslawski *et al*. 2016), 12 mg/mL α-syn monomer was dissolved in PBS and incubated at 37°C and 900 rpm for 5 h. After incubation, the sample was centrifuged, and the supernatant was applied to a size exclusion chromatography (SEC) column (Hiload 26/600 Superdex 200 pg) to separate the monomers from the oligomers (Figure 2B). Analysis of these fractions by SDS-PAGE under denaturing conditions showed the expected profile of monomeric and high molecular weight (HMW) bands, suggesting that the oligomer preparations contained a mixture of SDS-resistant and SDS-sensitive oligomers. An alternative explanation could be that the observed monomers were released from the ends/surfaces of the oligomers in the presence of SDS. The HMW species (with a molecular weight distribution of up to 1 MDa) could be visualized at the top of the resolving portion of the gel (Figure 2C). As expected, the monomers that were separated using SEC, it appeard in the gel around 15 kDa (Figure 2D), which was consistent with the expected MW of α-syn of 14461 Da (Figure 2E). The samples were analyzed by CD spectroscopy to ensure that each of the preparations exhibited the expected secondary structure signatures of oligomers and monomers. Oligomers exhibited a CD spectrum with a broad minimum peak centered at 219 nm, indicating the presence of mixed secondary structure contents dominated by β-sheet structures (Figure 2F). Monomers possessed a peak with the minimum at 198 nm consistent with their predominantly disordered structures (Figure 2F).

Next, we performed EM studies on monomer and oligomer preparations. Because of their small size (~14 kDa), the monomers are not visible by electron microscopy (Figure 2G). In contrast, the EM of the oligomer preparations showed heterogeneous morphologies consisting of annular porelike structures and spherical and rectangular tubular-like shaped particles (Figure 2H) (Lashuel *et al*. 2002) with a mean width of approximately 10 nm (Figure 2I). To prepare the fibrils, we followed the protocol described in Kumar et al., 2020. In brief, the lyophilized α-syn were dissolved in PBS to a final concentration of ~300 μM and incubated with shaking at 37 °C for five days at 1000 rpm. Next, we used our filtration protocol (Kumar *et al*. 2020a) to remove any remaining monomers and oligomers from the fibril preparations. The fibrils are enriched in β-sheet structures, as evidenced by the minimum peak at 221 nm in the CD spectrum shown in Figure 2J and the characteristic streaking pattern in the SDS-PAGE analysis, which confirmed the presence of SDS-resistant high molecular weight species of α-syn (Figure 2K). Ultrastructural analysis by EM revealed that these fibrils were polymorphic with fibril morphologies, including straight, twisted, or stacked fibrillar structures (Figure 2L), with a mean width of approximately 13 nm (Figure 2M).

### Stability of α-syn preparations

Since these preparations were to be characterized in different labs, we investigated their stability to ensure that they would not change their properties due to the shipping and storage conditions. Therefore, we subjected the oligomers to several cycles of freezing and thawing by snap-freezing them 3-4 times followed by room temperature thawing and incubation at temperatures of 4°C and 37°C for 2 days (Figure 2N). Interestingly, we found that the morphological distribution of the oligomers was not significantly altered when the oligomers were subjected to up to 3-4 freeze-thaw cycles or incubated at 4°C for 2 days (Figure 2O and 2I). However, the oligomeric mean width was slightly reduced by approximately 6% after incubation at 37°C (Figure 2O), which could be due to the release of α-syn monomers from the oligomeric structures.

We also tested the stability of the oligomer preparations using the digital ELISA assay based on the differences in the level of detection of oligomers at known concentrations under different solution conditions. Oligomers at 20 and 200 pg/mL were incubated at three different temperatures (2-8°C, +20-25°C or 36-38°C) for different durations (2, 4 and 24 h). In parallel, the effect of freezing and thawing (F/T) the samples 1x, 3x, and 5x was also tested. These samples were analyzed by the SIMOA sandwich assay using the A17183B antibody, which was captured on beads. The oligomer preparations are stable at 4°C for up to 24 hours and at 22°C for up to 4 hours. At temperature (37°C) for long incubation time (24 h), there was a significant decrease in the signal. When subjected to freeze/thaw cycles, a decrease in the signal was observed at lower (20 pg/mL) but not at higher concentrations (100 pg/mL) of oligomers. This evaluation did not include any optimization regarding the formulation of the oligomeric α-synuclein to ensure optimal stability in the follow-up experiments shown in Figure 4. However, these data, as well as the results of the EM analysis did not indicate any significant changes in the stability of the oligomers that could influence the interpretation of the results from the experiments performed in this study.

### Profiling the immunoreactivity of antibodies to different α-syn species by immunoblotting

Prior to our immunoblot experiments, we ensured equal amount of protein loading on the nitrocellulose membranes using a combination of Ponceau S staining (for slot blot analysis), Coommassie staining and silver nitrate staining (for Western blot analysis) (Supplementary figure 1). To assess the specificity of the antibodies, we first performed slot blot analysis of all the antibodies listed in Table 2 using pure preparations of α-syn monomers, oligomers, and fibrils under non-denaturing conditions (Figure 3A). Among the 18 antibodies tested (Figure 3B), 16 were reported in the literature to be conformation- or aggregation-state (oligomers or fibrils) specific; see Table 2 (Kovacs *et al*. 2012; Vaikath *et al*. 2015; Covell *et al*. 2017; van Diggelen *et al*. 2019). The remaining two antibodies, Syn 211 (which recognizes an epitope in the C-terminus region spanning residues 121-125) (Giasson *et al*. 2000) and SYN-1 (which recognizes an epitope in the NAC region spanning residues 91-99 of α-syn) (Perrin *et al*. 2003), are sequence-specific and recognized all three species. Surprisingly, we found that none of the antibodies tested (Table 2) had specific immunoreactivity towards only one particular species of α-syn (either the monomer, oligomers or fibrils). All 16 reported conformation-specific antibodies detected both oligomers and fibrils. Interestingly, among these, 1) the antibody clone 5G4 showed exceptional immunoreactivity towards oligomers and fibrils in a concentration-dependent manner (increased dose dependency) and almost no immunoreactivity towards monomers at both concentrations tested; 2) the antibodies SYNO3 and A17183E showed stronger immunoreactivity towards oligomers and fibrils but very weak (only at high concentrations) or no immunoreactivity towards monomers. Except for these three antibodies, the rest of the antibodies fell into one of the following three categories: i) antibodies that recognized oligomers and fibrils (even at low concentrations) with higher specificity than monomers (clones 26F1, SYNO2, and A17183B; ii) antibodies that recognized oligomers and fibrils in a concentration-dependent manner (high concentration → stronger detection) but also showed weak immunoreactivity towards monomers at high concentrations (clones 24H6, A17183A, SYNO4, 7015, 26B10, and A17183G); and 3) antibodies that were non-specific and recognized all three forms of α-syn (clones 9029, 12C6, ASyO5, and MJFR-14) (Figure 3A; summarized in Table 5).

**Figure 3:**
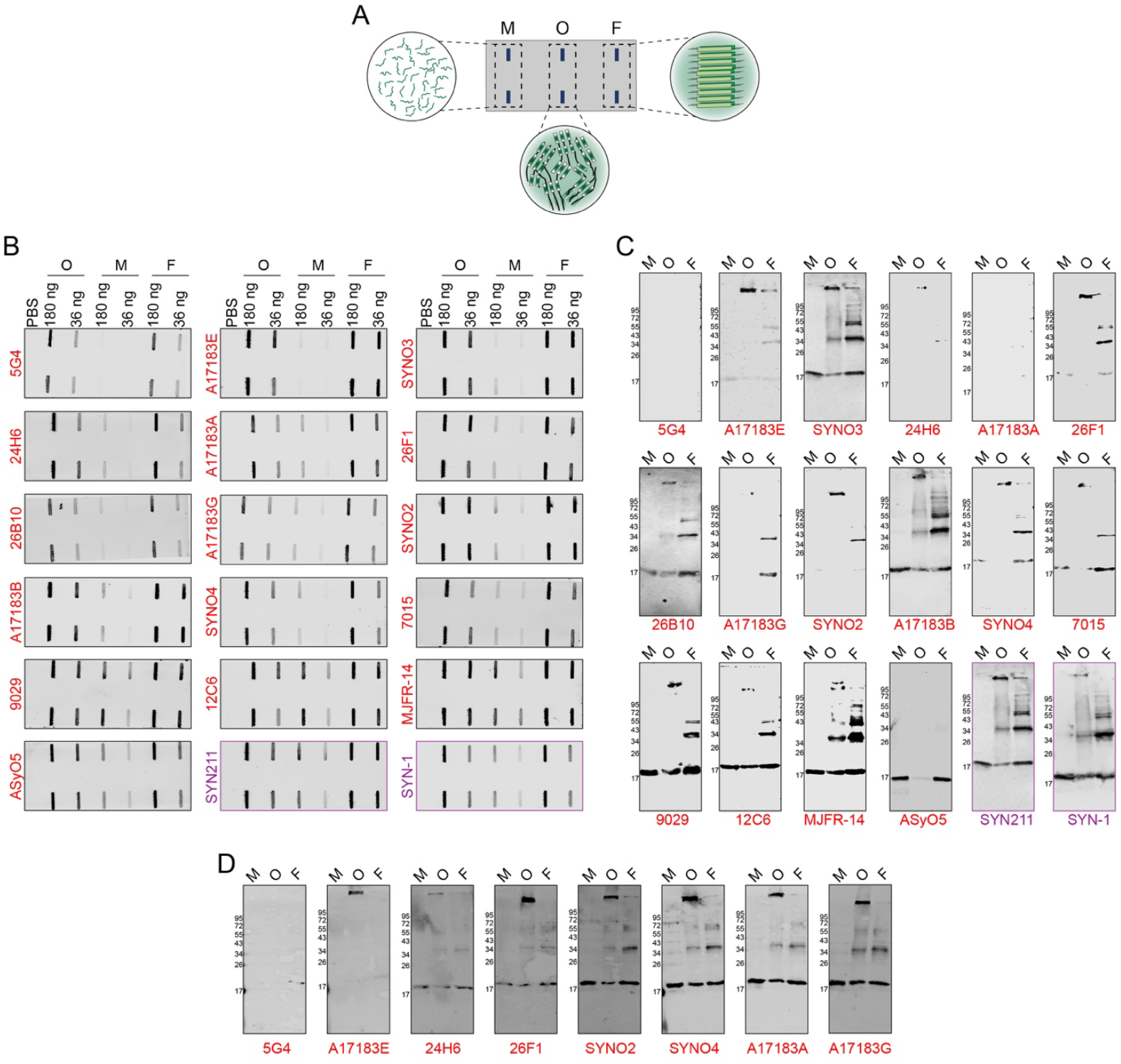
*In vitro* binding analysis of antibodies against α-syn monomers, oligomers and fibrils using slot blots and Western blots. (A) A schematic illustration of slot blot showing the blotting with different α-syn samples on the nitrocellulose membrane. (B) Slot blot analysis of the immunoreactivity of α-syn antibodies against α-syn monomers (M), β-sheet rich oligomers (O), and fibrils (F) under native conditions, spotted in duplicates at two different concentrations:180 ng and 36 ng. (C and D) Assessment of the immunoreactivity of antibodies against SDS and heat treated α-syn samples loaded at concentration of 36 ng (C) and 180 ng (D).

Given that many of these antibodies are also commonly used to assess the presence or formation of α-syn aggregates in cellular and animal models of synucleinopathies or in postmortem brain tissues using Western blot analysis, we assessed their immunoreactivity toward α-syn monomers, oligomers, and fibrils using this technique (Figure 3C). Although the samples are mixed and boiled in Laemmeli buffer which contains SDS prior to loading into the SDS-PAGE gels, it is not clear whether this treatment is sufficient to denature all the α-syn aggregates, i.e. the conformational state of the various α-syn species detected by Western blot remains undefined.

As expected, the sequence-specific antibodies SYN211 and SYN-1 showed stronger immunoreactivity towards monomers, oligomers and the high molecular weight bands in the fibrillar samples (Giasson *et al*. 2000; Perrin *et al*. 2003). This is consistent with the fact that the epitopes of these antibodies are outside of the domains that form the cores of oligomers and fibrils. Interestingly, antibodies such as 5G4, A17183E, 24H6, 26F1, A17183A, A17183G, SYNO2 and SYNO4 showed no or very weak immunoreactivity towards any of the three α-syn species when 36 ng the samples was used, suggesting that these antibodies recognize a native conformation that is lost upon treatment with SDS. Specifically, 5G4 and A17183A did not detect any of the α-syn bands. The 24H6 antibody reacted weakly only with the oligomeric band (at the top of the resolving gel). A17183E and 26F1 weakly detected monomers along with the detection of HMW bands. In contrast, several antibodies, including SYNO3, A17183B, 26B10, 9029, 12C6, AsyO5, MJFR-14 and 7015, which were reported to be oligomer/aggregate-specific, showed crossreactivity and detected SDS-denatured monomers, SDS-resistant oligomers and HMW bands in the fibrillar samples without any preference for one form of α-syn.

Given that some antibodies (5G4, A17183E, 24H6, 26F1, A17183A, A17183G, SYNO2 and SYNO4) had weak or no reactivity against α-syn at 36 ng, we repeated the Western blot analysis using a high concentration (180 ng) of α-syn. Interestingly, we observed similar results for 5G4 and A17183E (Figure 3D and Table 6), whereas 26F1 and 24H6 showed concentration-dependent reactivity to α-syn species (high concentration → stronger detection). However, the antibodies A17183A, A17183G, SYNO2 and SYNO4 which showed minimal detection at 36 ng (Figure 3C) displayed strong reactivity to α-syn bands at 180 ng (Figure 3D). Taken together, these results suggest that all the antibodies tested here do not preferentially detect one particular α-syn species (consistent with the slot blot analysis). Furthermore, Western blot analysis showed that many of the reported conformational- and aggregate-specific antibodies detected SDS-resistent HMW α-syn species and monomeric in SDS-PAGE gels (summarized in Table 6). These findings highlight the limitation of using selected antibodies to profile α-syn species by Western blot and underscore the critical importance of using multiple antibodies to capture the diversity of α-syn species.

Next, we assessed the antibody specificity towards α-syn monomers, oligomers and fibrils using a sandwich ELISA assay. We employed this assay for the detection of antibody specificity against low picogram concentrations of α-syn species under soluble conditions. The assay format utilizes the covalent capture of conformation-specific antibodies by a microsphere (Figure 4A). The three α-syn species described above were used as analytes at two concentrations (100 and 1000 pg/mL). A pan-synuclein antibody was included as a pairing antibody with the oligomer-specific antibody. When monomeric α-syn was used as the analyte, 12 of the 15 antibodies yielded lower than signal-to-noise S/N = 2, which is used as a threshold (Figure 4B). Three antibodies (clones 12C6, 7015 and 9029) showed S/N values higher than 2. In terms of immunoreactivity against oligomeric α-syn, 14 antibodies yielded S/N values higher than 2, although four antibodies (A17183E, 26F1, 24H6, and 26B10) possessed S/N values close to 2 in the presence of low concentrations of oligomers (100 pg/mL), but the immunoreactivity was enhanced at high concentrations (1000 pg/mL). The antibody 5G4 yielded the lowest S/N value of 1 at both concentrations (Figure 4C). When using fibrillar α-syn as an analyte, 14 antibodies resulted in S/N values higher than 2, while 26F1 showed immunoreactivity in the presence of high concentrations (1000 pg/mL) but had lower than borderline immunoreactivity toward 100 pg/mL of fibrils (Figure 4D). These experiments revealed that several antibodies reacted with monomeric α-syn, including 12C6 and 9029, whereas 7015 was borderline reactive. Those antibodies with strong immunoreactivity toward oligomers (10 antibodies: A17183A, A17183B, A17183G, SYNO2, SYNO3, SYNO4, 12C6, 7015, 9029 and MJFR14) also had strong immunoreactivity toward fibrils. The antibody 26F1 showed concentration-dependent immunoreactivity toward oligomers and fibrils (high concentration → stronger immunoreactivity), while the antibodies A17183E, 24H6, 26B10 and 5G4 showed greater fibril specificity. No antibodies could be identified that were solely oligomer-specific with no immunoreactivity toward α-synuclein monomers or fibrils, as observed by the slot blot analysis (Figure 3B) (summarized in Table 5).

**Figure 4:**
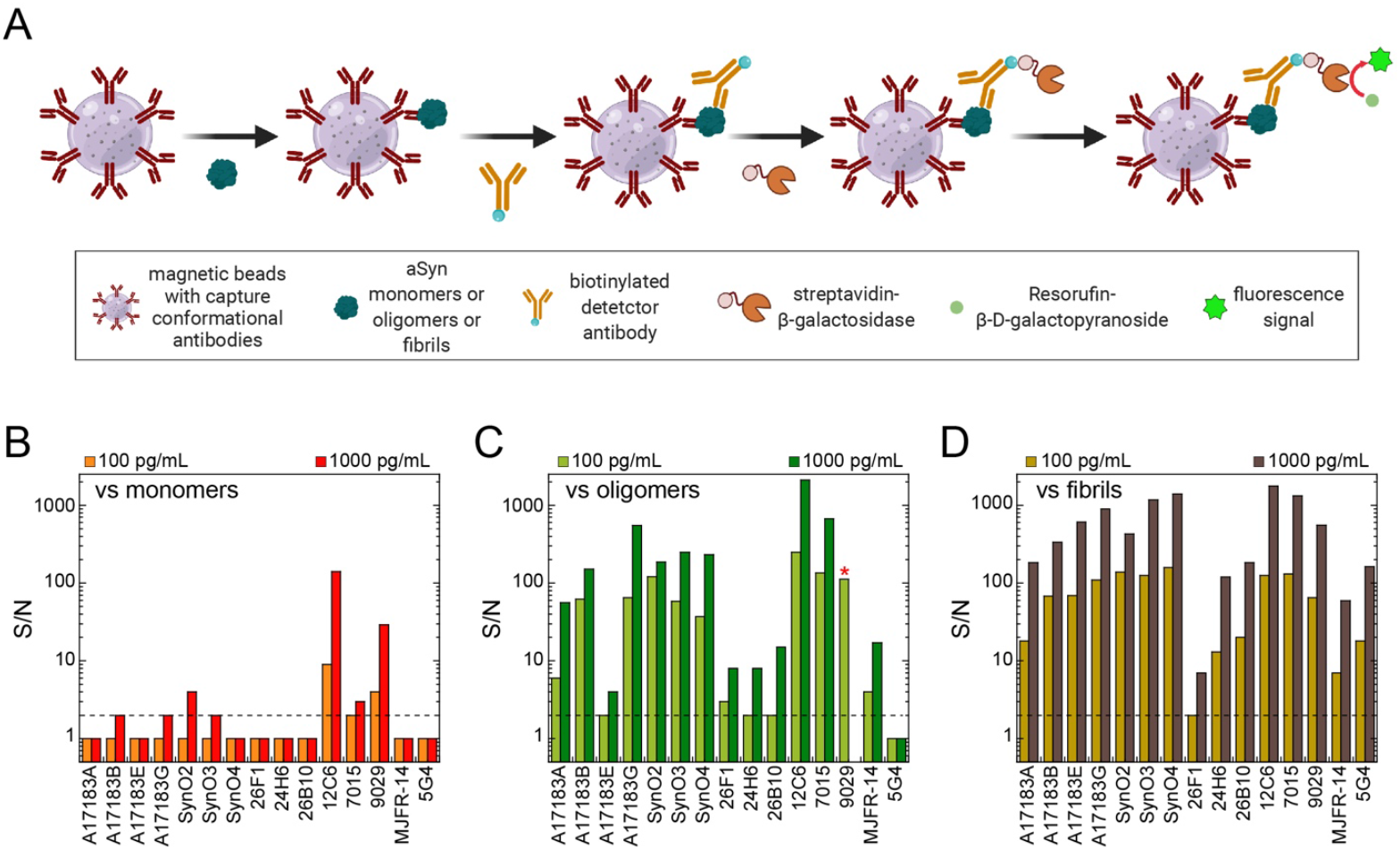
ELISA Simoa assay of antibodies against monomers, oligomers and fibrils. A) An illustration of ELISA Simoa assay and its experimental steps. The 15 α-syn conformation specific antibodies were coupled to the beads as capture monoclonal antibodies (X-axis), respectively. α-syn monomers (B), oligomers (C), and fibrils (D) were used as the analyte (100 and 1000 pg/mL). The detector antibody was a C-terminal monoclonal antibody (clone SYN211) with an epitope in the amino acid region from 121-125. The signal-to-noise values (S/N) are indicated on the Y-axis for the three α-syn forms. * indicates the availability of only 100 pg/mL oligomer data for antibody clone 9029 (C).

### Characterization of the specificity of the antibodies toward morphologically and structurally different forms of oligomeric α-syn

Since our knowledge of the morphological and conformational properties of native oligomers in the brain is unknown, we sought to further assess the specificity of the antibodies toward different preparations of oligomers that exhibited distinct structurally, chemical and morphologically properties. The use of these different oligomer preparations allowed us to test whether differences in the morphologies/structures of α-syn oligomers could influence the immunoreactivity or binding specificities of the antibodies. We employed dopamine (DA) (Figure 5A) and 4-hydroxy-2-nonenal (HNE) (Figure 5B) to prepare cross-linked human WT α-syn oligomers. Several studies have shown that the interaction of DA with α-syn promotes the formation of α-syn oligomers and influences α-syn aggregation propensity and neurotoxicity (Conway *et al*. 2001; Mor *et al*. 2017), raising the possibility of the presence of DA-modified α-syn oligomeric species in PD patient brains. Similarly, HNE, a physiological byproduct of lipid peroxidation, has been shown to play roles in oxidative stress responses and to alter the aggregation of α-syn in PD (Qin *et al*. 2007; Ingelsson 2016).

**Figure 5:**
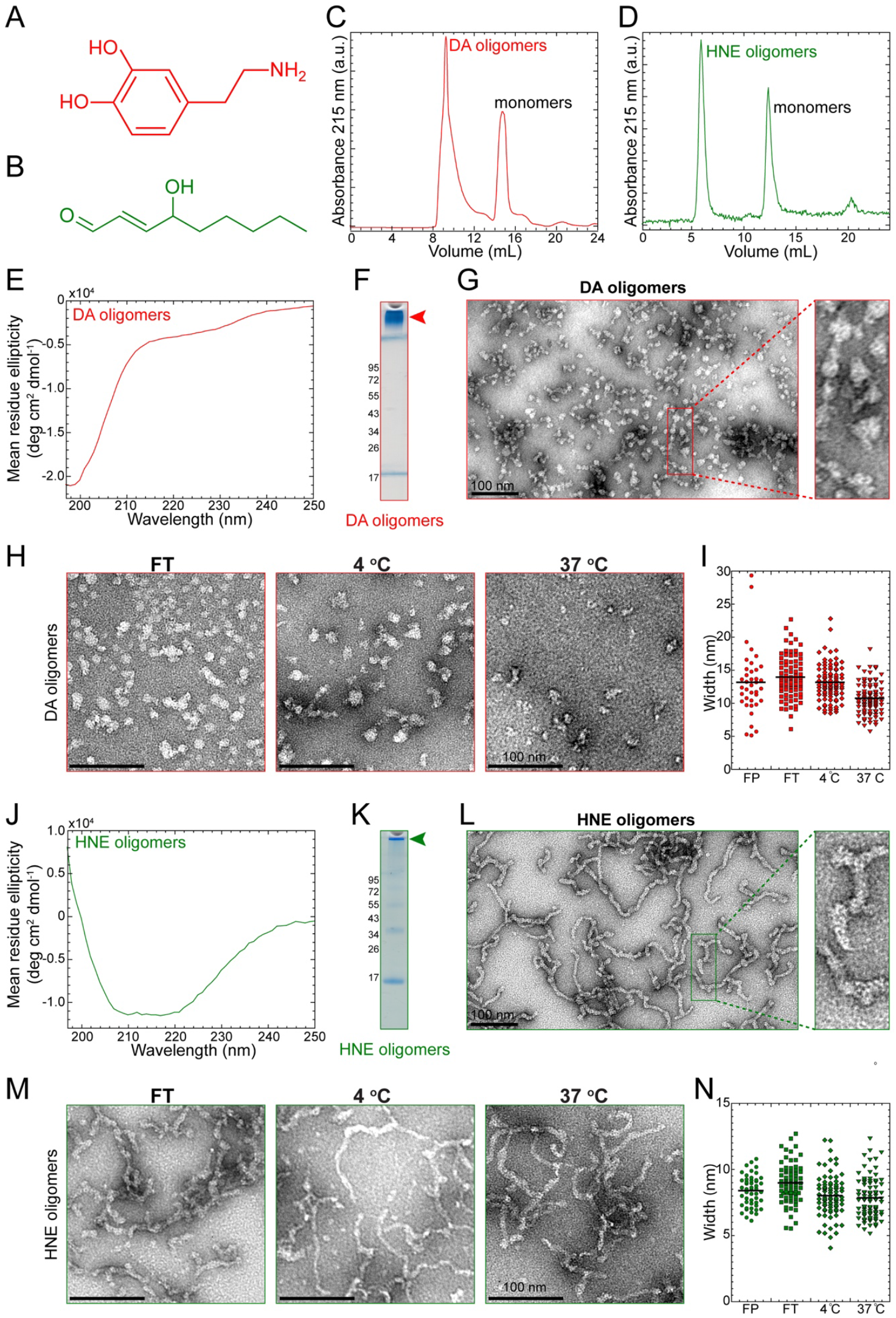
Preparation of dopamine- and HNE-induced oligomers and analysis of their stability. Chemical structures of dopamine (A) and HNE (B). SEC purification of DA-induced oligomers (C) and HNE-induced oligomers (D) separated from monomers. (E) CD spectra of DA-induced oligomers. (F) Coomassie staining of dopamine and DA-induced oligomers. (G) Negatively stained EM analysis performed on DA-induced oligomers. DA-induced oligomers showed mainly spherical, undefined morphologies. (H) Assesment of the stability of the DA-induced oligomers subjected to different temperature conditions incubation (4 °C and 37 °C) and and freeze-thawing as determined by analysis of their width distribution (I). (J) CD spectra of HNE-induced oligomers. (K) Coomassie staining of HNE-induced oligomers. (L) Negatively stained EM analysis performed on HNE-induced oligomers. (M) Assessment of the stability of HNE-induced oligomers incubation (4 °C and 37 °C) and and freeze-thawing as determined by analysis of their width distribution (N).

The DA- and HNE-induced oligomers (Figure 5 C and D) were prepared by the incubation of DA or HNE with α-syn, followed by the isolation of the oligomers using SEC as described above. Mass spectrometry analysis of the monomeric fractions from the SEC purifications (in both DA- and HNE-induced oligomer preparations; Figure 5C and 5D) showed masses that are higher than the expected mass of monomeric α-syn (14460 Da). In the samples where α-syn was co-incubated with dopamine, we observed an increase in mass by 65 Da (14525 Da; Supplementary figure 2A), which may correspond to the oxidation of the four methionine present in α-syn (4*16=64 Da). For the α-syn monomers isolated by SEC from the HNE-α-syn sample mixtures, we observed several peaks reflecting the addition of single or multiple modification of 156 Da each (14615 Da, 14771 Da, 14928 Da and 15084 Da; Supplementary figure 2B), corresponding to the formation of HNE-α-syn adducts. The DA-induced oligomers exhibited CD spectra with a minimum at 198 nm, revealing the presence of species with predominantly disordered conformations and little structure (Figure 5 E, Table 4). However, the HNE-induced oligomers showed a broad CD spectrum centered at 219 nm that is more similar to the CD spectrum of the oligomers (Figure 2F), indicating these oligomers are rich in β-sheet structure (Figure 5J, Table 4). Analysis of these fractions by SDS-PAGE analysis under denaturing conditions (Figure 5F: DA oligomers, Figure 4K: HNE-induced oligomers) showed a very similar gel profile for both types of oligomers, with the presence of HMW bands at the top of the resolving part of the gel and a light band of monomers at 15 kDa that may have been released from the oligomers in the presence of SDS.

EM ultrastructural analysis of the DA-induced oligomers showed the presence of oligomers with near-spherical morphologies of different shapes and sizes (Figure 5G), as previously shown (Conway *et al*. 2001; Cappai *et al*. 2005; Mahul-Mellier *et al*. 2015). These oligomers exhibited a mean width of approximately 13 nm (Figure 4I). However, the HNE-induced oligomers appeared to be more homogenous and displayed a curvilinear (chain-like) morphologies with a mean width of approximately 8 nm (Figure 5L and 5-N).

Next, we tested the stability of these oligomers by monitoring changes in their sizes and oligomeric morphologies as described above for the oligomers shown in Figure 2. Neither repeated cycles of freeze-thaw conditions or longer incubations at 4 °C seem to influence the morphologies of the DA oligomers. The number/density of the oligomeric particles on the EM grids was significantly reduced for the DA oligomeric sample incubated at 37 °C (Figure 5H), and a decrease in the mean width to approximately 10 nm was also observed (Figure 5I). Under identical conditions, the HNE oligomers did not show major changes in their morphologies or mean width (Figure 5M and 5N).

To investigate whether the antibodies in Table 2 show differential immunoreactivity to morphologically, chemically and conformationally distinct oligomer preparations under native conditions, we performed slot blot analysis using DA- and HNE-induced oligomers and monomers as a control (Figure 6A). The majority of the antibodies we tested detected DA- and HNE-induced oligomers and monomers irrespective of their morphological or structural differences (Figure 6B). Strikingly, among all the antibodies tested, 26F1 showed no immunoreactivity toward DA-induced oligomers, weak immunoreactivity toward monomers but very strong immunoreactivity toward HNE-induced oligomers, which is consistent with the data shown in Figure 3B. 26F1 was generated against oligomerized HNE-modified α-syn. The 5G4 antibody, which does not recognize α-syn monomers, also showed strong immunoreactivity toward HNE-induced oligomers but exhibited weak immunoreactivity toward DA-induced oligomers, which could be due to the presence of small population of structured DA oligomers (Table 4) or due to the weak affinity of 5G4 to the ensemble of DA-induced oligomers. As expected, the sequence-specific antibodies SYN211 and SYN-1 detected both DA- and HNE-induced oligomers and monomers.

**Figure 6:**
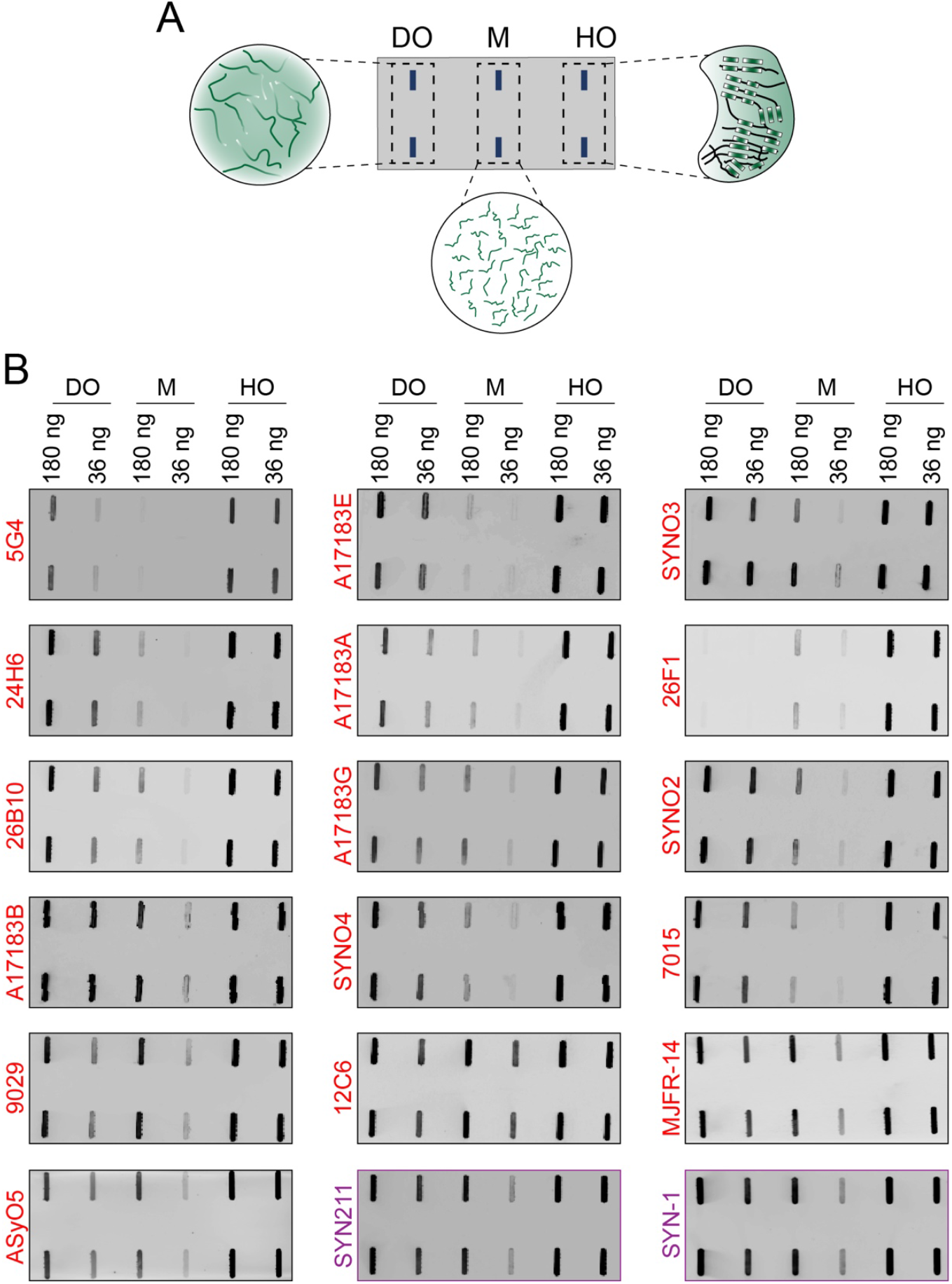
Slot blot analysis of antibodies against α-syn monomers and DA- and HNE-induced oligomers. (A) A scheme illustrating the slot blot analysis showing the different α-syn samples blotted in the nitrocellulose membrane. (B) Assessment of the immunoreactivity of antibodies by slot blot against dopamine-induced unstructured oligomers (DO), α-syn monomers (M) and HNE-induced oligomers (HO), and fibrils (F) under native conditions and spotted in duplicates at two different concentrations: 180 ng and 36 ng.

The antibodies that showed stronger immunoreactivity toward HNE-induced oligomers can be categorized further depending on their immunoreactivity toward DA-induced oligomers and monomers as follows: i) A17183E showed enhanced detection of DA- and HNE-induced oligomers compared to monomers; ii) 24H6, 9029, ASyO5, A17183A, 26B10, MJFR-14 and A17183G showed stronger immunoreactivity to HNE-induced oligomers (even at a low concentration of protein of 36 ng per spot) and concentration-dependent detection of DA-induced oligomers and monomers; iii) SYNO2, SYNO3, SYNO4, A17183B, 7015 showed enhanced detection of HNE- and DA-induced oligomers and concentration-dependent immunoreactivity toward monomers (weak binding at a low concentration of monomers, 36 ng of protein per spot). In contrast, 12C6 showed very strong nonspecific detection of DA- and HNE-induced oligomers as well as monomers irrespective of the concentration of proteins loaded in each spot in the slot blot analysis, consistent with the ELIA results.

Next, we investigated whether the concentration of antibodies may influence their immunoreactivities towards different α-syn species. To do this, we selected few antibodies that were shown to be aggregate specific (26F1, 5G4) and others that are nonspecific and recognize different α-syn forms (Syn211, SYN-1, 9029, SynO4, MJFR-14 and ASyO4). We reassessed their specificity over antibody concentrations ranging from 2 to 200 ng/mL) and observed similar results (Supplementary figure 3) as reported in (Figure 3B, Figure 6B) at all the antibody concentrations tested.

In summary, our immunoblotting studies (Figure 3, Figure 6 and Supplementary figure 3) demonstrate that none of the antibodies showed any preferential specificity toward one particular α-syn species, including monomers, oligomers and fibrils. However, we observed some exceptions: i) the antibody clone 26F1 did not show any immunoreactivity toward the largely unstructured DA-induced oligomers but was highly specific for β-sheet-enriched oligomers, HNE-induced oligomers and fibrils; ii) the antibody 5G4 showed weak immunoreactivity toward the largely unstructured DA-induced oligomers but stronger immunoreactivity towards β-sheet-enriched oligomers, HNE-induced oligomers and fibrils (summarized in Table 5). These differences in immunoreactivity emphasize the importance of using many antibodies in parallel rather than a single antibody for the identification of pathological oligomers in the brain given the heterogeneous nature and structural properties of such oligomers.

### Kinetics of the binding of α-syn monomers and oligomers to immobilized antibodies

To further characterize and validate the specificity of the antibodies, we assessed their binding affinity and kinetics to monomeric and oligomeric α-syn species using SPR (Figure 7A). The antibodies were immobilized on the SPR chip surface at low densities. Figure 7 and supplementary figure 4 show the SPR sensorgrams of selected antibodies (seven in total; six conformational and one sequence-specific) as a function of time as obtained by the successive injection of monomers or oligomers at concentrations ranging from 30 to 2000 nM. Sensorgram plots were fitted to extract the kinetic parameters, such as the binding affinity (KD) and association (k_a_) and dissociation (k_d_) rate constants of the binding between antibodies and monomer or oligomer complexes. The fitting of the plots was based on either a 1:1 binding model or by a global heterogeneous ligand binding model, which provides kinetic parameters for two binding sites. Collectively, all of the tested antibodies showed binding responses to both α-syn monomers and oligomers with varying binding affinities. Interestingly, many of the antibodies that were found to be highly specific for oligomers/fibrils (Figure 3B) also showed some binding to monomers. However, the binding affinities (KD) of the antibodies A17183A, SYNO4, and 26F1 reflected μM affinities toward monomers, which was consistent with the slot blot data (Figure 3B). The kinetic parameters obtained from the sensorgrams are summarized in Table 3.

**Figure 7:**
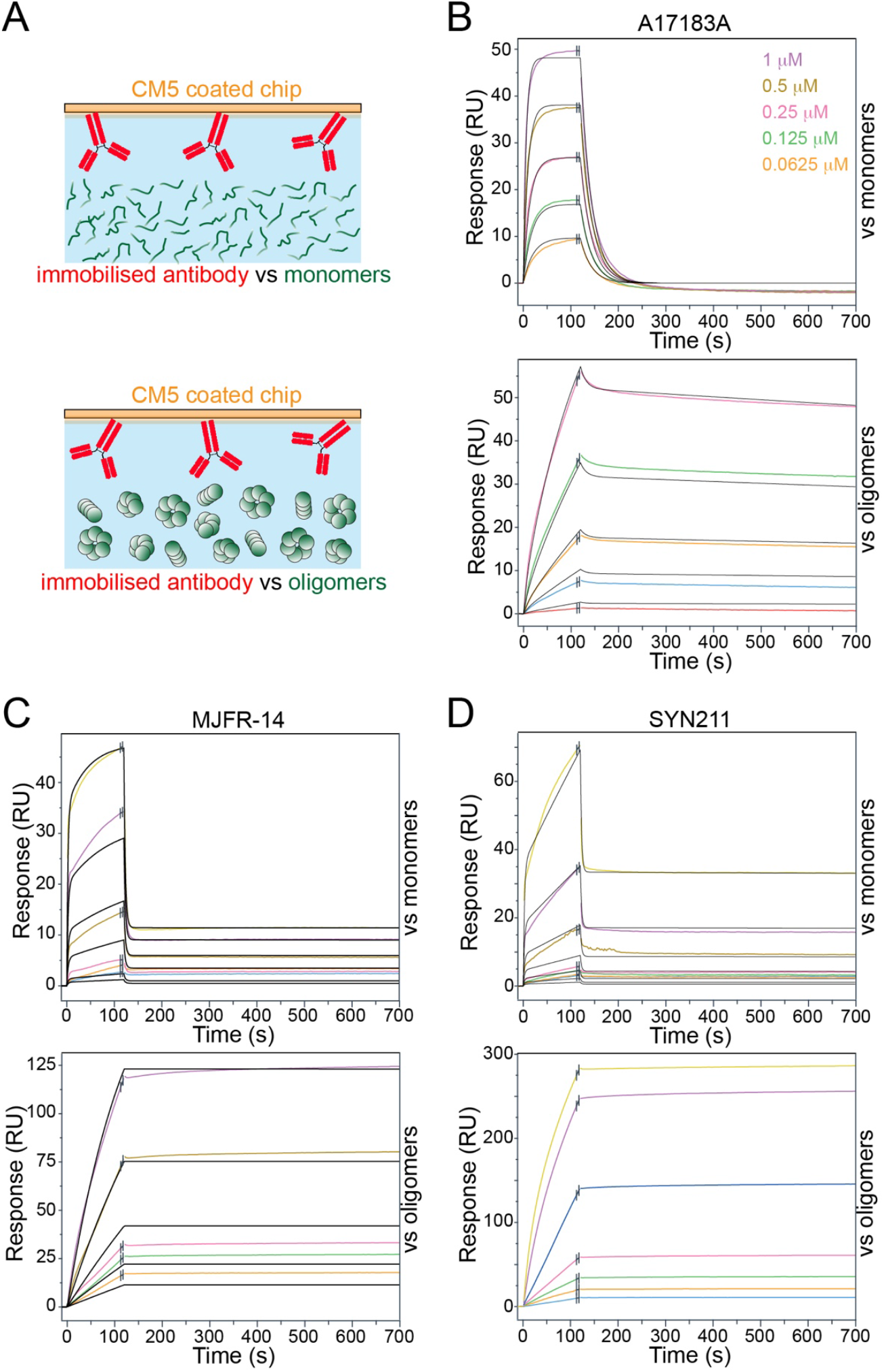
SPR-based kinetic analysis of different immobilized monoclonal antibodies (A17183A (B), MJFR-14 (C), and SYN211 (D)) binding to α-syn monomers (top) and oligomers (bottom) at 30, 60, 120, 250, 500, 1000 and 2000 nM concentrations. The antibodies were immobilized at a ligand density of approximately 3000-4000 RUs. The α-syn monomers or oligomers were injected for 2 min, followed by 5 min dissociation with injection of PBS buffer at a 30 μL/min flow rate. The sensorgrams are shown as colored lines representing varying concentrations of α-syn monomers or oligomers, and the fits are shown as black lines. The kinetic parameters obtained from the fitting are shown in Table 3.

**Table 3:**
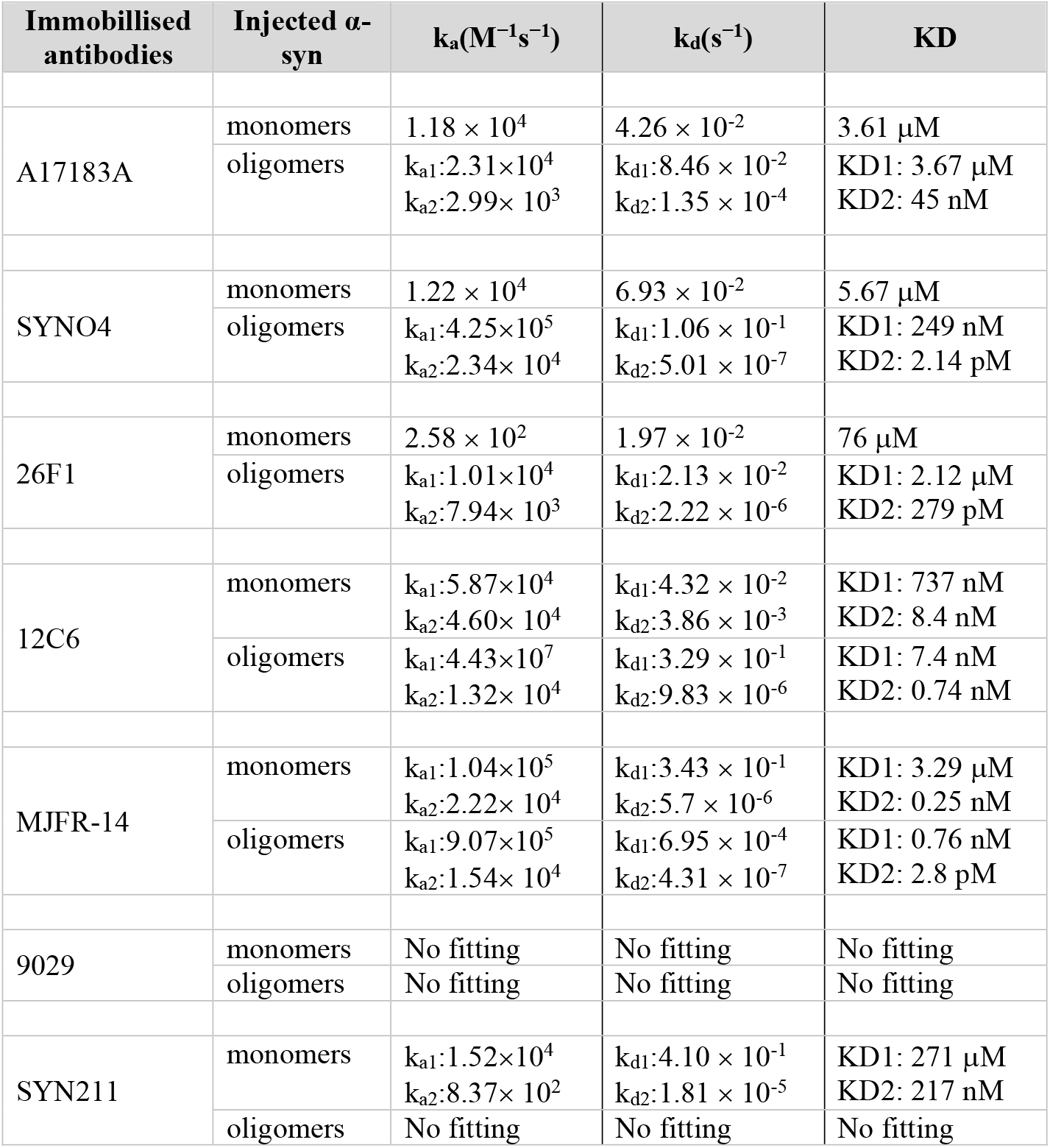
Kinetic parameters of the antibodies binding to α-syn monomers or oligomers

The shape of the dissociation portion of the SPR sensorgrams (after 120 seconds) provides clues about the binding affinity of the antibodies toward monomers or oligomers; a stronger affinity is reflected by a flatter slope of the dissociation curve (slower off-rate), but a weaker affinity is reflected by a steeper curve (faster off-rate). Figure 7 and supplementary figure 4 show the SPR sensorgrams of the antibodies upon titration of monomers and oligomers. Fitting with a 1:1 binding model was possible for the binding of a few antibodies to α-syn monomers. These antibodies showed weak binding affinities to monomers with KD values of of 3.61 μM (A17183A), 5.67 μM (SYNO4) and 76 μM (26F1). However, the same antibodies exhibited stronger binding affinities towards oligomers as evident by the fitting of the raw data which was possible only by heterogeneous binding model, suggesting two possible oligomer binding sites: A17183A (KD1: 3.67 μM, KD2: 45 nM), SYNO4 (KD1: 249 nM, KD2: 2.14 pM) and 26F1 (KD1: 2.12 μM, KD2: 279 pM) (Figure 7B and SI Figure 1A, Table 3). A stronger binding affinity for oligomers by antibodies A17183A, SYNO4 and 26F1, is in agreement with our slot blot analyses (Figure 3B), confirming that these antibodies bind with greater specificity to oligomers than monomers. 12C6 and 9029 showed stronger binding toward oligomers (Figure 7C and Supplementary figure 4, Table 3), but also exhibited good binding to monomers. This is consisitent with the fact that these antibodies showed binding to monomers by slot blots and ELISA (Figure 4B) and detected α-syn under denaturing conditions. However, it was not possible to fit the data and calculate the kinetic rate constants and binding affinities for 9029. Both antibodies showed a slow dissociation of monomers as evidenced by the fact that the dissociation portion of the sensorgrams (after 120 seconds) returns slowly to the baseline response units (RU, y-axis) but not as rapidly as seen with A17183A, 26F1 and SynO4. These observations suggest that the antibodies 9029 and 12C6 possessed a stronger affinity to monomers, in comparison to A17183A, 26F1 and SynO4, which exhibit weak binding affinity to monomers.

The MJFR-14 antibody also showed preference for binding oligomers, but still showed binding to monomers. The SPR sensograms suggested two distinct kinetics events on the dissociation portion with a rapid drop (after 120 seconds) indicating a major fraction of monomers dissociating fastly because of the weak affinity to MJFR-14 (KD1: 3.29 μM). However, the RU values did not return to baseline, suggesting that MJFR-14 might have a second binding to another conformation of monomers (KD2: 0.25 nM) or binds strongly to a small population of α-syn aggregates which might have formed from the monomers during the SPR experimental timeframe. MJFR-14 also showed two binding kinetics against oligomers, but, both exhibited very stronger affinitiy (KD1: 0.76 nM and KD2: 2.8 pM). As expected, SYN211, which recognizes all three forms of α-syn, showed a stronger affinity for monomers (Figure 7D). The fitting of SYN211 binding to oligomers was not possible but indicated stronger binding to oligomers, as reflected by the shape of the curves.

## Discussion

Increasing evidence supports the hypothesis that different forms of α-syn aggregates (e.g. fibrils and oligomers) play an important role in the pathogenesis of PD. Testing this hypothesis requires the development of therapeutic drugs or antibodies that target the different species and assays that enable the accurate assessment of changes in their levels during disease progression and in response to therapies. Although there are several biochemical, structural and imaging-based approaches for the direct and indirect visualization and characterization of α-syn fibrils (Shahmoradian *et al*. 2019; Lashuel 2020), detection of nonfibrillar oligomeric α-syn species in cells or postmortem brain tissues remains challenging. The existing methods and techniques, such as Western blotting and proximity ligation assays, provide indications of the presence of oligomers but not information about their size, conformation and/or morphology. The instability and low abundances of native oligomers make the isolation and characterization of their structural properties using NMR and Cryo-EM very challenging. Due to these challenges, researchers in the field have resorted to the development of conformation- or aggregate-specific antibodies.

One of the major untested assumptions about conformation-specific antibodies is that they are capable of capturing the structural and morphological diversity of α-syn aggregates *in vivo* or to target specific α-syn aggregates. However, these assumptions are rarely experimentally tested. Therefore, there is a need to develop protocols and pipelines that enable systematic characterization of antibodies using a well-characterized and validated set of α-syn reagents representing, to the extent possible, the diversity of α-syn in vivo. Towards this goal, we developed such a validation pipeline and used it to evaluate the binding specificity of 18 α-syn antibodies, 16 of which were reported to be conformation- or aggregation state-specific (Table 2). First, the antibodies were screened against disordered monomers and preparations of β-sheet rich α-syn oligomers and fibrils. This enabled us to test the specificity of the antibodies to monomers, oligomers and fibrils. To further assess whether antibody binding was indeed driven by conformational specificity or avidity, the antibodies were screened against three preparations of α-syn oligomers with distinct biochemical, conformational and morphological properties (disordered oligomers and oligomers with different β-sheet and secondary structure contents) (Table 4). In parallel, we also tested the specificity and binding affinities of various α-syn antibodies by SIMOA assays and SPR, respectively.

**Table 4:** Morphologies and structural properties of oligomers used in this study

| Oligomers | Morphologies | Width/diameter | Secondary structure composition (%) |  |  |
| --- | --- | --- | --- | --- | --- |
| | | | α-helix | β-strand | irregular |
| Oligomers | annular pore,<br>spherical,<br>tubular | 6-14 nm | 27 | 38 | 32 |
| DA<br>oligomers | near-spherical,<br>globular | 5-20 nm | 11 | 11 | 78 |
| HNE<br>oligomers | curvi-linear | 5-11 nm | 27 | 26 | 49 |

**Table 5:** Immunoreactivity and binding specificity of antibodies against native structures of α-syn species by different techniques

| Immunogen | Antibodies | Slot blot analysis |  |  |  |  | ELISA assay |  |  | SPR |  |
| --- | --- | --- | --- | --- | --- | --- | --- | --- | --- | --- | --- |
|  |  | M | O | DO | HO | F | M | O | F | M | O |
| oligomers | 26F1 | + | +++ | -- | +++ | +++ | -- | ++ | ++ | * | *** |
|  | 12C6 | +++ | +++ | +++ | +++ | +++ | +++ | +++ | +++ | *** | *** |
|  | 24H6 | + | +++ | +++ | +++ | +++ | -- | ++ | +++ | Ø | Ø |
|  | 26B10 | + | ++ | +++ | +++ | ++ | -- | ++ | +++ | Ø | Ø |
| fibrils | 7015 | ++ | +++ | +++ | +++ | +++ | + | +++ | +++ | Ø | Ø |
|  | 9029 | ++ | +++ | +++ | +++ | +++ | ++ | +++ | +++ | Ø | Ø |
|  | SYNO2 | + | +++ | +++ | +++ | +++ | + | +++ | +++ | Ø | Ø |
|  | SYNO3 | + | +++ | +++ | +++ | +++ | + | +++ | +++ | Ø | Ø |
|  | SYNO4 | + | +++ | +++ | +++ | +++ | -- | +++ | +++ | * | *** |
| recombinant α-syn aggregates | A17183A | + | +++ | +++ | +++ | +++ | -- | +++ | +++ | * | *** |
|  | A171183B | + | +++ | +++ | +++ | +++ | + | +++ | +++ | Ø | Ø |
|  | A17183E | + | +++ | +++ | +++ | +++ | -- | + | +++ | Ø | Ø |
|  | A17183G | + | ++ | +++ | +++ | +++ | + | +++ | +++ | Ø | Ø |
| a.a. (44-57) | 5G4 | + | +++ | ++ | +++ | +++ | -- | -- | +++ | Ø | Ø |
| a.a. (1-140) | SYN211 | +++ | +++ | +++ | +++ | +++ | Ø | Ø | Ø | *** | Ø |
| a.a. (15-123) | SYN-1 | +++ | +++ | +++ | +++ | +++ | Ø | Ø | Ø | Ø | Ø |
| a.a. (111-125) | ASyO5 | ++ | +++ | ++ | +++ | +++ | Ø | Ø | Ø | Ø | Ø |
| α-syn filaments | MJFR-14 | +++ | +++ | +++ | +++ | +++ | -- | ++ | +++ | * | *** |

M: monomers; O: oligomers; DO: DA induced oligomers; HO: HNE induced oligomers; F: fibrils.
-- : no detection; + : faint detection; ++ : medium detection; +++ : strong detection; Ø : data not available
\*\*\* : strong affinity; \* : weak affinity

**Table 6:**
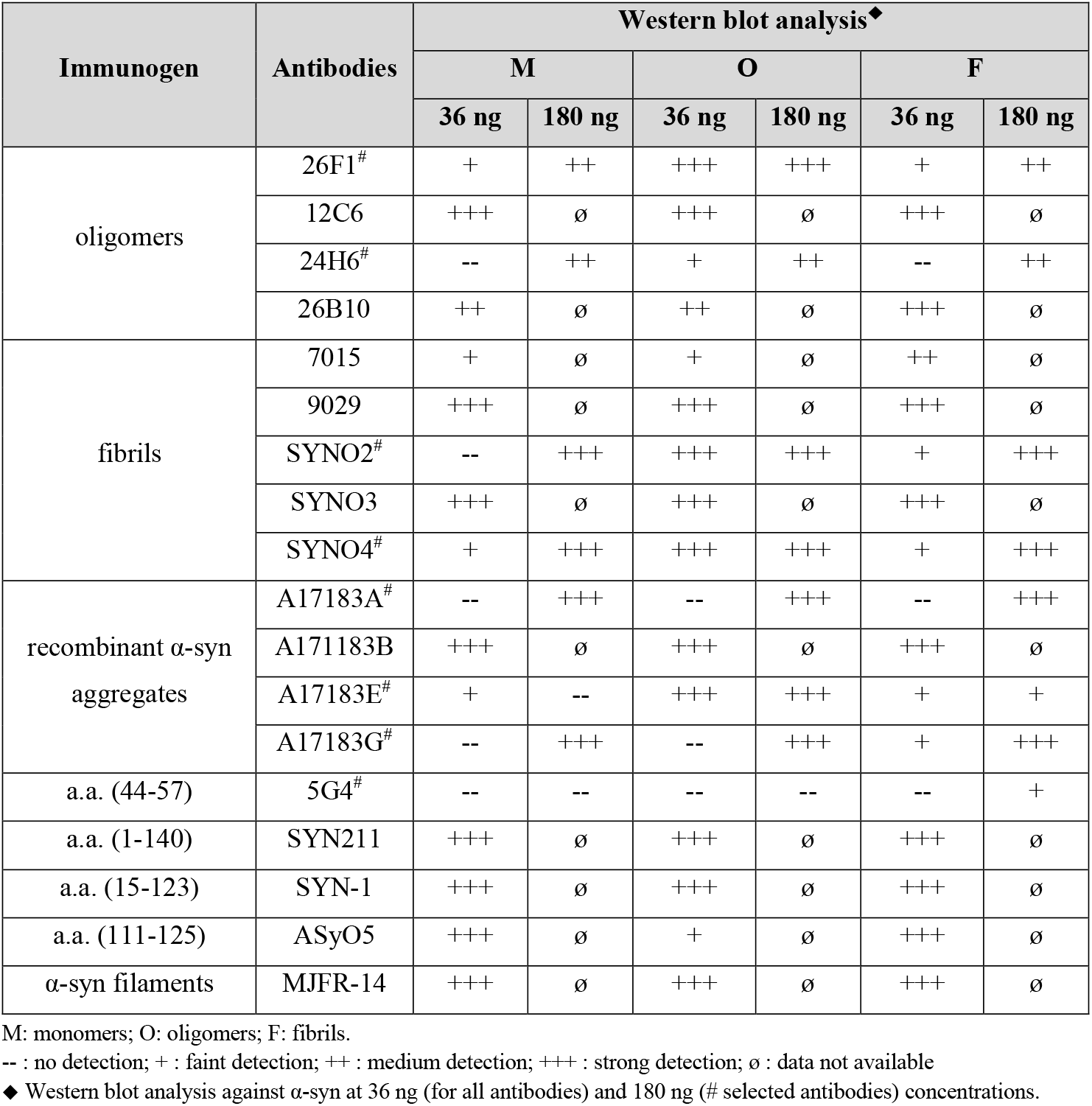
Immunoreactivity of antibodies against SDS and heat treated α-syn samples by Western blot analysis

### None of the antibodies specifically detect either monomers, oligomers or fibrillar α-syn species

Surprisingly, we found that none of the antibodies tested in our study had unique specific immunoreactivity toward one particular α-syn species (monomers, oligomers or fibrils). All 16 reported conformational-specific antibodies detected both unmodified β-sheet rich oligomers and fibrils, demonstrating that they could not differentiate between oligomers and fibrils and are not specific for a particular conformation or α-syn aggregation state.

In an attempt to assess the specificity of the antibodies against a diverse set of oligomers, we also produced oligomeric preparations (DA and HNE oligomers) possessing structurally, chemically and morphologically distinct properties (Figure 5). The dopamine-induced oligomers were predominantly disordered, whereas the HNE-induced oligomers were rich in β-sheet structure. The immunoreactivity toward these oligomers was compared to that toward the unmodified oligomers, which are enriched in β-sheet structures (Figure 3 and Figure 6). Despite the similarity of the CD signatures of HNE-induced and unmodified α-syn oligomers, the two types of oligomers exhibited distinct morphological features (Figure 2 and Figure 5). The structural and morphological diversity of the different oligomer preparations provided a unique opportunity to assess the specificity of the conformational-specific antibodies.

As expected, the Syn 211 and SYN-1 antibodies detected all three forms of α-syn species (monomers, oligomers and fibrils) as well as the DA- and HNE oligomers in both slot blots and Western blots (Figure 3) (Giasson *et al*. 2000; Perrin *et al*. 2003). This could be attributed to the fact that their epitopes are located in regions that do not constitute the core of α-syn oligomers or fibrils (Paslawski *et al*. 2014b; Li *et al*. 2018; Guerrero-Ferreira *et al*. 2019) and are thus likely to be exposed in both aggregation states. Our results are consistent with previous studies in which Syn 211 was shown to detect monomers and HNE-induced oligomers (van Diggelen *et al*. 2019) and the detection of monomers and fibrils by SYN-1 (Vaikath *et al*. 2015; Weihofen *et al*. 2019). Interestingly, the antibody clone 5G4 showed increased immunoreactivity with high conformational specificity for all forms of α-syn aggregates but showed very weak immunoreactivity toward monomers. We can not rule out the possibility that the binding to the DA oligomers could arise from the presence of partial β-sheet structure or small population of oligomers with β-sheet structure in these preparations, as suggested by the analysis of the CD data (Table 4). Previous studies reported that 5G4 detects widespread and distinct α-syn-induced pathology in the cortical and brain stem brain regions in postmortem synucleinopathic brain tissues (Kovacs *et al*. 2012) but only weakly detects monomeric bands in brain homogenate samples from Lewy body dementia patients. Furthermore, van Diggelen et al. found that 5G4 antibody detected HNE-induced oligomers and showed no immunoreactivity toward monomers (van Diggelen *et al*. 2019).

However, several of our observations with a number of antibodies were not consistent with previously published reports or data provided by the manufacturers of the antibodies. Previous reports indicated that the antibody MJFR-14 is a conformational-specific antibody that detects filamentous α-syn aggregates (Sampson *et al*. 2016; Elfarrash *et al*. 2019; Kawahata *et al*. 2019), but not monomeric form of the protein. Martinez *et al*. reported that MJFR-14 is a conformation-specific with enhanced immunoreactivity towards filaments but not to the denatured filaments or monomers by dot blot analysis (Martinez T. N. 2016; abcamCat.No.ab209538). Until the publication of a preprint version of this report in bioRxiv (Kumar *et al*. 2020b), the antibody was sold by (abcamCat.No.ab209538) as “Anti-Alpha-synuclein filament antibody [MJFR-14-6-4-2] - Conformation-Specific”. Furthermore, data obtained using Luminex assay demonstrated an increased specificity of MJFR-14 antibody towards α-syn oligomers compared to monomers and filaments (Martinez T. N. 2016) at low ng concentrations). Interestingly, it was previously shown that MJFR-14 exhibits weaker binding to monomers, which could be eliminated by preabsorbing the antibodies with recombinant α-syn (Martinez T. N. 2016). MJFR-14 has also been described as an oligomer-specific antibody. For example, Lassen *et al*. reported that MJFR-14 is highly specific for oligomers but not to monomers (Lassen *et al*. 2018). Direct comparison to fibrils was not performed in this study. In line with these evidence, our ELISA Simoa assay showed that MJFR-14 does not bind to monomers at low picogram concentrations (Figure 4B) and although it binds to both to fibrils and oligomers, it exhibits preferential binding to fibrils compared to oligomers (Figure 4C and D). In addition, our slot blot analysis (Figure 3B and Figure 6B) showed stronger and similar immunoreactivity towards oligomers and fibrils but a weaker immunoreactivity to monomers at 36 ng concentrations. Collectively, our studies show that MJFR-14 shows high immunoreactivity toward all aggregated forms of α-syn, including unmodified oligomers, DA- and HNE-induced oligomers and fibrils, suggesting that this antibody is neither fibril- or oligomer-specific. Furthermore, MFJR-14 binds to both β-sheet rich and disordered α-syn oligomers. Altogether, our findings confirm previous reports suggesting preferential binding of MFJR-14 to aggregate forms of α-syn, but also show that it still binds to monomers in a concentration-dependent manner (Figure 3, Figure 4, Figure 6, Figure 7C and Supplementary figure 3) and recognizes α-syn monomeric bands in SDS-PAGE gels (Figure 3C). These observations combined with MFJR-14 strong immunoreactivity towards disordered oligomers (DA oligomers) suggest that its preferential binding to oligomers and fibrils could be driven by avidity rather than by its conformational specificity.

Similarly, Vaikath et al. reported that SYNO2, SYNO3, and SYNO4 can bind to α-syn oligomers and fibrils but not monomers (Vaikath *et al*. 2015). However, in our study, SYNO2 and SYNO3 recognized strongly oligomers and fibrils and detected α-syn monomers in a concentrationdependent manner (higher concentration of monomer → better immunoreactivity) (Figure 3 and Figure 6). Again, all three antibodies recognized equally β-sheet rich (oligomers and HNE oligomers and disordered oligomers (DA oligomers), suggesting that their specificity to α-syn aggregates could be driven by avidity rather than conformational specificity. Interestingly, all three antibodies SYNO2, SYNO3, and SYNO4 recognized monomeric α-syn bands in SDS-PAGE gels (Figure 3C and D).

ASyO5 is another commercial antibody (Agrisera: AS13 2718) that has been reported to exclusively detect oligomers, but not monomers or fibrils (using dot blot, (AgriseraCat.No.AS132718). However, in our hand and using the α-syn samples described above, we found that ASyO5 antibody binds non-specifically to α-syn monomers, different types of oligomers and fibrils (Figure 3B, Figure 6B and Supplementary figure 3).

Among the 18 antibodies in Table 2, the binding affinity and kinetics of only three antibodies (SYNO2, SYNO3, and SYNO4) against α-syn fibrils but not monomers or oligomers have been described in the literature or in the material provided by the manufacturer (Vaikath *et al*. 2015). Most importantly, we could not find any comparative SPR or binding studies using well-characterized preparations of α-syn monomers or different types of oligomers.

The comparison of the kinetics data and binding affinities of various antibodies for monomers and oligomers showed a significant degree of variation in the values of k_a_, k_d_, and KD (Table 3). Antibody clones A17183A, 26F1, and SYNO4 showed high binding affinities for oligomers and weak binding affinities for monomers (Figure 7, Supplementary Figure 4 and Table 3). This suggests that these antibodies are highly conformationally specific, which is in agreement with the slot blot and Western blot data (Figure 3B and C). Interestingly, the A17183A antibody showed stronger immunoreactivity to unstructured DA oligomers (Figure 6), which may hint that the binding is, perhaps, driven by avidity rather than affinity. However, the 26F1 antibody showed specificity for β-sheet-enriched oligomers but not for unstructured DA oligomers, suggesting that it could be truly conformationally specific. The very weak affinity of 26F1 with a KD of 76 μM for monomers is also in line with all our analyses, including the ELISA analysis, suggesting that 26F1 is highly conformationally specific for β-sheet-enriched α-syn aggregates.

Other antibodies, such as 7015, 12C6, 9029 showed strong binding and immunoreactivities for monomers, oligomers and fibrils, although 7015 showed higher binding and immunoreactivity to oligomers and fibrils. By ELISA, the antibodies 12C6, 7015, and 9029 exhibited high immunoreactivity towards monomers at low pg concentrations (Figure 4B; 100 and 1000 pg/mL concentration of monomers used). This is consistent with both the slot blot and Western blot analyses, where these antibodies showed high immunoreactivity to monomers (Figure 3B) and detection of monomeric α-syn bands in denaturing gels (Figure 3C).

### Limitations of our study

One major limitation of our work is that while we used diverse and well-characterized α-syn preparations of monomers, oligomers (different types), and fibrils to screen the antibodies, it remains unclear to what extent these species occur in the brain. That being said, we hypothesized that screening using a diverse set of species instead of using one specific type of α-syn oligomer is the best that we can do to approximate the complexity of α-syn species *in vivo*. The second limitation is that α-syn is subjected to different modifications *in vivo*, while all our protein standards were generated from unmodified recombinant α-syn. However, it is important to note that while we know a great deal about the different types of PTMs that occur in LBs and LNs and α-syn aggregates, very little is known about the PTM patterns of α-syn oligomers *in vivo*. Further studies are needed to address this knowledge gap. Finally, our studies focused on exploring the structural diversity of oligomers but not that of fibrils. We recognize this limitation and plan to address it in future studies.

## Conclusions

Herein, we used multiple techniques to assess the α-syn species specificity of several commonly used conformational-specific and aggregation state α-syn antibodies. This was achieved using well-characterized preparation of α-syn monomers, fibrils and different preparation of oligomers of distinct structural and biochemical properties. Our results demonstrated that i) no antibodies could be identified that were solely monomer-specific, oligomer-specific or fibril specific; ii) all the antibodies that recognized α-syn oligomers also recognized α-syn fibrils and some recognized all three species (oligomers, fibrils and monomers); iii) the antibody clone 26F1 is the only antibody that was shown to be highly specific for β-sheet-enriched oligomers, it detects oligomers and HNE-induced oligomers and fibrils but not for unstructured DA-induced oligomers and structurally disordered monomers; All other antibodies recognized both structured (β-sheet enriched) and disordered oligomers, suggesting that their specificity could be driven by avidity rather than conformational specificity iv) the antibody clone 5G4 showed increased immunoreactivity toward β-sheet-enriched oligomers, HNE-induced oligomers and fibrils, and unstructured DA-induced oligomers and almost no immunoreactivity toward monomers; iv) antibodies clones A17183A, A17183E, SYNO4 preferentially detected all three types of oligomers and fibrils but reacted very weakly toward monomers; v) the majority of the other antibodies (such as 9029, 12C6, ASyO5, SYN-1, and SYN211) exhibited immunoreactivity towards all α-syn species under the conditions tested here. MJFR-14 shows more specificity to aggregated forms of α-syn by ELISA, but showed higher immunoreactivity to monomers by slot blot and Western blotanalyses. Although we failed to identify antibodies that target a single specific form of α-syn, i.e. monomers, oligomers or fibrils, our results show that it is possible to develop antibodies that target β-sheet rich α-syn oligomers and fibrils or oligomers and fibrils of diverse conformational properties. Such antibodies could represent more reliable tools for measuring the total levels of aggregated α-syn. Finally, our findings show that it is unlikely that any of the existing oligomer-specific immunoassays are capable of providing an accurate assessment of the levels of α-syn oligomers or capturing the diversity of α-syn by Western blots and possible in tissues. Therefore, we propose that these oligomer assays should be reassessed for their ability to distinguish between α-syn oligomers and fibrils and that interpretation of previous and future studies should take into account the specificity and limitation of the antibodies used. Future studies aimed at deciphering the role of different α-syn species in the pathogenesis of PD should be carried out using multiple antibodies that have been characterized using α-syn multiple calibrants that capture, to the extent possible, the diversity of α-syn species in the brain. A similar approach can be applied to facilitate the development of accurate assays to assess target engagement of therapeutic antibodies.

## Authors’ contributions

HAL, HV, ES and conceived and conceptualised the study. HAL STK, SJ, ES and designed the experiments. STK and SJ performed all the experiments except ELISA. CF carried out the ELISA assay. STK, SJ, ES and HAL analysed the data. STK, SJ and HAL wrote the manuscript, with inputs from CF and ES.

## Acknowledgments

We are grateful to Dr. Ron Gill (senior scientist, Xsensio SA, Switzerland) for his critical review, helpful discussions and comments on the SPR data. We thank the CIME facility (EPFL) for the use of their electron microscopy facility and Dr. Kelvin Lau from PTPSP (Protein Production and Structure Core Facility, EPFL) for their tremendous support and use of their SPR and CD instruments. We also thank Guus Scheefhals (Syngle Therapeutics B.V.), Dr. Ramanath Hegde, Dr. Pedro Magalhães, Dr. Anne-Laure Mahul-Mellier, Dr. Salvatore Novello, Firat Melek Altay and Ahmed Sadek (EPFL, Lausanne) for their critical review and feedback on the paper. We also thank Dr. Rajashekhar Kolla for his help with the drawing of chemical structures of dopamine and HNE. We also thank Anass Chiki for providing the purified α-syn used for the preparation of DA- and HNE-induced oligomers, and Jonathan Ricci for his assistance with the initial setup of immunoblot studies. Antibodies with clones 7015 and 9029 were kindly provided by Dr. Kelvin Luk, Prof. John Trojanowski and Prof. Virginia M. Lee (UPenn, USA). The generation of antibodies 24H6, 12C6, 26F1 and 26B10 was supported by a grant from the Michael J. Fox Foundation (DOSAB). This project was funded by the École Polytechnique Fédérale de Lausanne and the Michael J Fox Foundation.

## Conflict of interest disclosure

Prof. Hilal A. Lashuel is the founder and chief scientific officer of ND BioSciences, Epalinges, Switzerland.

## Materials and methods

### Recombinant overexpression and purification of human WT α-syn

Recombinant overexpression and purification of human WT α-syn was performed as described previously (Fauvet *et al*. 2012) with slight modifications. pT7-7 plasmids encoding human WT α-syn were used for transformation in BL21 (DE3) *E-Coli* cells on an ampicillin agar plate. A single colony was transferred to 400 mL of Luria broth (LB) medium containing 100 μg/mL ampicillin (AppliChem, A0839) (small-scale culture) and incubated overnight at 37 °C and 180 rpm. On the next day, the pre-culture was used to inoculate 3-6 liters of LB medium having 100 μg/mL ampicillin (large-scale culture). Upon A_600_ approaching 0.4 to 0.6, α-syn protein expression was induced by the addition of 1 mM 1-thio-β-d-galactopyranoside (AppliChem, A1008) and the cells were further incubated at 37 °C and 180 rpm for four hours. This incubation step was followed by harvesting cells by centrifugation at 4000 rpm using JLA 8.1000 rotor (Beckman Coulter, Bear, CA) for 30 minutes at 5 °C. The harvested pellets were stored at −20 °C until the next step. The cell lysis was performed by dissolving the bacterial pellet in buffer A (20 mM Tris–HCl, pH 7.5) containing protease inhibitors (1 mM EDTA (Sigma-Aldrich, 11873580001) and 1 mM PMSF (Applichem, A0999) followed by ultrasonication (VibraCell VCX130, Sonics, Newtown, CT) time: 5 min; cycle: 30 sec ON, 30 sec OFF; amplitude 70%. After lysis, centrifugation at 12000rpm and 4°C for 30minutes was performed to collect the supernatant. This supernatant was collected in 50 mL Falcon tube and placed in boiling water (100 °C) for about 15 minutes. This solution was subjected to another round of centrifugation at 12000 rpm and 4 °C for 30 minutes. The supernatant obtained at this step was filtered through 0.45 μm filters and injected into a sample loop connected to HiPrep Q FF 16/10 (GE healthcare, 28936543). The supernatant was injected at 2 mL/min and eluted using buffer B (20mM Tris-HCl, 1M NaCl, pH 7.5) from gradient 0 to 70% at 3 mL/min. All fractions were analyzed by SDS-PAGE, and the fractions containing pure α-syn were pooled and concentrated using a 30 kDa molecular weight cut-off (MWCO) filters (MERCK, UFC903008) at 4 °C. The retentate was collected and dialyzed using 12-14 kDa MWCO Spectrapor dialysis membrane (Spectrum Labs, 9.206 67) against deionized water at 4 °C overnight to remove salts. Dialyzed solution was collected, snap-frozen, and lyophilized.

### Preparation of WT α-syn oligomers

To generate monomer and fibril free α-syn oligomeric preparations, 60 mg of lyophilized recombinant α-syn protein was dissolved in 5 mL PBS (10 mM disodium hydrogen phosphate, 2 mM potassium dihydrogen phosphate, 137 mM NaCl and 2.7 mM potassium chloride, pH 7.4) (final concentration: 12mg/mL) containing 5 μL Benzonase (final concentration: 1 μL benzonase/mL (MERCK, 71205-3). After dissolving, the solution is filtered through 0.22 μm filters (MERCK, SLGP033RS) and transferred to five low-protein binding 1.5 mL tubes, each containing 1 mL solution. These tubes were incubated in a shaking incubator at 37 °C and 900 rpm for five hours. The samples were centrifuged at 12000*g* for 10 min at 4 °C to remove any insoluble α-syn aggregates. 5 mL of supernatant was loaded into a sample loop of the chromatography setup. This sample was run through Hiload 26/600 Superdex 200pg (GE Healthcare, 28-9893-36) column equilibrated with PBS and eluted as 2.5 mL fractions at a flow rate of 1 mL/min. The elution of protein was monitored by UV absorbance at 280nm. Different fractions were visualized by SDS-PAGE analysis, and fractions of interest (oligomer) corresponding to the void volume peak were aliquoted (500 μL), snap-frozen, and stored at −20 °C.

### Preparation of DA-induced oligomers

DA-induced oligomers were prepared as described previously (Mahul-Mellier *et al*. 2015). Briefly, the recombinant α-syn protein was dissolved in 20mM Tris and 100mM NaCl to have a final concentration of 140 μM (pH 7.4). After dissolving, the protein solution was filtered through a 100 kDa filter (MERCK, MRCFOR100). The filtrate was transferred to a low-protein binding tube and 20 equivalents of dopamine (final concentration: 2.8 mM) (Sigma-Aldrich, H8502) was added. This tube was covered with aluminum foil and incubated in a shaking incubator at 37 °C and 200 rpm for five days. The sample was centrifuged at 12000*g* for 10 min at 4 °C to remove any insoluble α-syn aggregates. The supernatant was loaded into a sample loop of the chromatography setup. This sample was then run through Superdex 200 Increase 10/300 GL (GE healthcare, 28990944) column equilibrated with PBS and eluted as 0.5 mL fractions at a flow rate of 0.4 mL/min. The elution of protein was monitored by UV absorbance at 280 nm. The SEC fractions were analyzed by SDS-PAGE analysis, and fractions of interest (oligomer) were collected and stored at 4 °C.

### Preparation of HNE-induced oligomers

HNE-induced oligomers were prepared as described previously (Näsström *et al*. 2011a). Briefly, the recombinant α-syn protein was dissolved in 20 mM Tris and 100 mM NaCl to have a final concentration of 140 μM (pH 7.4). After dissolving, the protein solution was filtered through 100 kDa filter (MERCK, MRCFOR100). The filtrate was transferred to a low-protein binding tube and 30 equivalents of HNE (Cayman Chem, 32100) (final concentration: 4.2mM) was added. This tube was incubated in an incubator at 37°C under quiescent conditions for 18 hours. Following incubation, the sample was centrifuged at 12000g for 10 min at 4 °C to remove any insoluble α-syn aggregates. The supernatant was loaded into a sample loop of the chromatography setup. This sample was run through Superdex 200 Increase 10/300 GL (GE healthcare, 28990944) column equilibrated with PBS and eluted as 0.5 mL fractions at a flow rate of 0.4 ml/min. The elution of protein was monitored by UV absorbance at 280nm. The SEC fractions were analyzed by SDS-PAGE analysis, and fractions of interest (oligomer) were collected and stored at 4 °C.

### Protein concentration estimation

The concentration of α-syn samples such as monomers, oligomers, and fibrils were estimated using BCA assay and amino acid analysis. For BCA assay, microplate measurements were carried out using BCA protein assay reagents (Pierce, catalog number: 23227). Briefly, triplicates of known concentrations (from 10 μg/mL-1000 μg/mL) of bovine serum albumin (concentration standard) and an equal volume of α-syn samples were pipetted into microplate wells. To which, 200 μL of BCA working reagent was added and incubated at 37 °C for 30 minutes. Absorbance at 562 nm was measured using a Tecan plate reader. Using BCA assay based concentration estimation as standards, known concentrations (2-3 μg) of α-syn samples were pipetted into a conical insert, flash-frozen, and lyophilized. The dried form of α-syn samples was shipped to Functional Genomic Center Zurich for subjecting for amino acid analysis (AAA) for absolute quantification of α-syn samples concentrations.

### Preparation of WT α-syn fibrils

WT α-syn fibrils were prepared as described in (Mahul-Mellier *et al*. 2020). Briefly, 4 mg of lyophilized recombinant α-syn was dissolved in 50 mM Tris, and 150 mM NaCl and pH was adjusted to 7.5. The solution is filtered through 0.2 μm filters (MERCK, SLGP033RS), and the filtrate is transferred to black screw cap tubes. This tube was incubated in a shaking incubator at 37 °C and 1000 rpm for five days. After five days, the formation of fibrils was assessed by TEM and SDS-PAGE, followed by Coomassie staining as described in (Mahul-Mellier *et al*. 2018).

### Characterization of oligomers

#### SDS-PAGE analysis

Human WT α-syn monomers, unmodified WT, DA-induced, and HNE-induced oligomers were run on 15% polyacrylamide gels, and Coomassie blue staining was performed as described previously (Kumar *et al*. 2020a). Before carrying out Western blot analysis, equal loading on the gel was also confirmed using Coomassie staining or silver staining (Invitrogen, LC6100) as per manufacturers protocol.

#### TEM analysis

TEM analysis of protein samples were performed as described previously (Kumar *et al*. 2020a). Briefly, 5 μL protein samples were placed on glow discharged Formvar and carbon-coated 200 mesh-containing copper EM grids. After about a minute, the samples were carefully blotted using filter paper and air-dried for 30 s. These grids were washed three times with water and followed by staining with 0.7% (w/v) uranyl formate solution. TEM images were acquired by Tecnai Spirit BioTWIN electron microscope, and image analysis was performed by ImageJ software as described previously (Kumar *et al*. 2020a).

#### Far-UV circular dichroism (CD) spectroscopy

Approximately 150 μL of protein samples were loaded onto 1 mm path length quartz cuvette, and CD spectra were obtained on Chirascan spectropolarimeter (Applied Photophysics) with the following parameters as described in (Kumar *et al*. 2020a). Temperature: 20 °C; wavelength range: 198 to 250 nm; data pitch: 0.2 nm; bandwidth: 1 nm; scanning speed: 50 nm/min; digital integration time: 2 s. The final CD spectra was a binomial approximation on an average of 10 repeat measurements. The secondary structural content of the oligomers was estimated using the online CD analysis tool known as CAPITO (Wiedemann *et al*. 2013).

#### Mass spectrometry analysis

Mass spectrometry (MS) analysis of proteins were performed by liquid chromatography-mass spectrometry (LC-MS) on the LTQ system (Thermo Scientific, San Jose, CA). Before analysis, proteins were desalted online by reversed-phase chromatography on a Poroshell 300SB C3 column (1.0×75mm, 5um, Agilent Technologies, Santa Clara, CA, on the LTQ system). 10 uL protein samples were injected on the column at a flow rate of 300 uL/min and were eluted from 5 % to 95 % of solvent B against solvent A, linear gradient. The solvent composition was, Solvent A: 0.1% formic acid in ultra-pure water; solvent B: 0.1% formic acid in acetonitrile. MagTran software (Amgen Inc., Thousand Oaks, CA) was used for charge state deconvolution and MS analysis.

#### Temperature stability analysis of oligomers

Human WT α-syn unmodified WT, DA-induced and HNE-induced oligomers were tested for their stability at different temperature conditions. Morphological characteristics were assessed by TEM under conditions such as freeze-thaw cycles, storage at 4°C or 37°C for 5 days.

### Binding characterization of α-syn species to different antibodies

#### Slot blot analysis

Nitrocellulose membranes (Amersham, 10600001) were spotted with 5 μL and 1 μL samples volumes corresponding to 180 ng and 36 ng of α-syn proteins in duplicates (monomers, oligomers, DA-induced oligomers, HNE-induced oligomers and fibrils) per sample spot. The membranes were blocked for 1 hour with Odyssey blocking buffer (LiCoR, 927-40000), and then incubated overnight with different primary antibodies (Table 2) diluted in PBST at 2 μg/mL concentration for all the antibodies except SYN-1 antibody at 1 μg/mL concentration (Figure 3 and 6). Varying concentrations of primary antibodies (200, 20 and 2 ng/mL) were used in the Supplementary figure 3. The membranes were washed three times, with 0.1% PBS-Tween (3×10minutes) and incubated with IRdye conjugated secondary antibodies (1:7500) (Table 1) for 1 hour at RT. Thereafter, the membranes were washed three times, with 0.1% PBS-Tween (3×10minutes). The visualization was performed by fluorescence using Odyssey CLx from LiCor. Equal loading of protein samples on the membrane was confirmed using Ponceau S (MERCK, P3504) staining (2% Ponceau S (w/v) in 5% acetic acid).

#### Western blot analysis

Approximately 36 ng and 180 ng of proteins (monomers or oligomers or fibrils) were loaded onto 15 % SDS-PAGE gels (prior to loading samples were boiled at 95°C for 10 minutes) and run at 180 V for 1 hour in running buffer (25 mM Tris, 192 mM Glycine, 0.1% SDS, pH 8.3). Gels were transferred onto nitrocellulose membranes (Amersham, 10600001) at 25 V, 0.5 A, and 45 minutes using Trans-Blot Turbo (Bio-Rad, 170-4155). The membranes were blocked for 1 hour with Odyssey blocking buffer (LiCoR, 927-40000), and then incubated overnight with different primary antibodies (Table 2) diluted in PBST at 2 μg/mL concentration for all the antibodies except SYN-1 antibody at 1 μg/mL concentration. The specificity of MJFR-14 was also assessed at 2 ng/mL against 180 ng of α-syn monomer, oligomers and fibrils (Supplementary Figure 3B). The membranes were washed three times, with 0.1% PBS-Tween (3×10minutes) and incubated with IR dye conjugated secondary antibodies (1:7500) (Table 1) for 1 hour at RT. After that, the membranes were washed three times, with 0.1% PBS-Tween (3×10 minutes). The visualization was performed by fluorescence imaging using Odyssey CLx from LiCor.

#### Determination of antibody affinities by surface plasmon resonance (SPR, BIACORE)

SPR data were collected on a Biacore 8K device (GE Healthcare). Antibodies were immobilized on a CM5 biosensor chip (GE Healthcare) at 10-20 μg/mL concentration in 10 mM acetate solution (GE Healthcare) at pH 4.5 to reach a final surface ligand density of around 2000-4000 response units (RUs). In short, the whole immobilization procedure using solutions of 1-ethyl-3-(3-dimethyl aminopropyl) carbodiimide (EDC) and N-hydroxy succinimide (NHS) mixture, antibody sample and ethanolamine, was carried out at a flow rate of 10 μl/min into the flow cells of the Biacore chip. Firstly, the carboxyl groups on the sensorchip surface were activated by injecting 200 μL of 1:1 (v/v) mixture of EDC/NHS (included in the amine coupling kit, Cytiva life sciences) into both flow cells 1 and 2 and followed by the injection of antibodies overflow cell 2 for 180 s. The remaining activated groups in both the flow cells were blocked by injecting 129 μL of 1 M ethanolamine-HCl pH 8.5. The sensor chip coated with antibodies were equilibrated with PBS buffer before the initiation of the binding assays. Serial dilutions of analytes such as α-syn monomers or oligomers (oligomers) at a concentration ranging between 2 μM to 0.015 μM in PBS buffer were injected into both flow cells at a flow rate of 30 uL/min at 25 °C. Each sample cycle has the contact time (association phase) of 120 seconds and followed by a dissociation time of 600 seconds. After every injection cycle, surface regeneration of the Biacore chips was performed using 10 mM glycine (pH 3.0). The obtained data were processed and analyzed using Biacore 8K evaluation software for the calculation of the binding kinetics (association rate constant (Ka) and dissociation rate constant (Kd)) and binding affinity (KD). The fitting of the data was based on either 1:1 binding model (mostly for the monomers) or heterogeneous ligand binding model (for oligomers) using global kinetic fitting unless otherwise noted.

#### Antibody characterization and stability analysis of oligomers by digital sandwich ELISA using Simoa technology

The Quanterix Simoa platform is a highly sensitive platform that allows detection at the sub pg/mL concentration range (Rissin *et al*. 2010). Attempts to measure for example neurofilament light into plasma matrix were successful by Simoa technology as described before (Kuhle *et al*. 2016). Kuhle et al compared three immunoassays for neurofilament light chain measurements into blood. The analytical sensitivity was 78 pg/mL and 0.62 pg/ml for the conventional ELISA and the Simoa based assay, respectively. The presence of conformational synuclein forms (oligomers and fibrils) into body fluids like CSF is expected to be low abundant making the Simoa platform the preferred technology platform for our experimental work.

A sandwich ELISA Simoa immunoassay was used to assess the immunoreactivity of different α-syn antibodies towards α-synuclein forms such as monomers, oligomers, and fibrils. To prepare the conjugated beads, paramagnetic carboxylated particles/beads were activated for 15 minutes at 4 °C using 0.05 mg/mL of 1-ethyl-3-[3-dimethylaminopropyl]carbodiimide hydrochloride (EDAC) (ThermoScientific, Cat N°A35391) added to 1,40 ×10^9^ beads/mL. The beads were washed using a magnetic separator, and 0.1 mg/mL of the oligomeric α-synuclein specific monoclonal capture antibody was added. After 2 hours of incubation on a mixer-shaker at 4 °C, the conjugation reaction was blocked (Quanterix blocking buffer, Cat N°101356) for 30 minutes at room temperature. The conjugated beads were washed and stored at 4 °C. The biotinylated detector antibody (SYN211) was used with an antibody/biotin ratio of 64. The assay was performed on the fully automated Quanterix Simoa HD-1 with a 2-step protocol. The undiluted samples were tested, which required 300 μL volume of samples without accounting for dead volume (duplicate testing). The calibrator diluent consists of 1xPBS with 0.1% milk, 0.1% Tween. The first incubation step of the sample with the beads was 60 minutes. After washing, the second incubation step with streptavidin-β-galactosidase (Quanterix, Cat N° 103397) was 5 minutes. Prior to reading Resorufin-β-D-galactopyranoside (Quanterix, Cat N° 101736) was added. The resulting fluorescence signal is captured and translated into an AEB value (Average Enzymes per Bead) that is proportional to the analyte concentration in the measured sample.

## Supporting information

**Supplementary figure 1:**
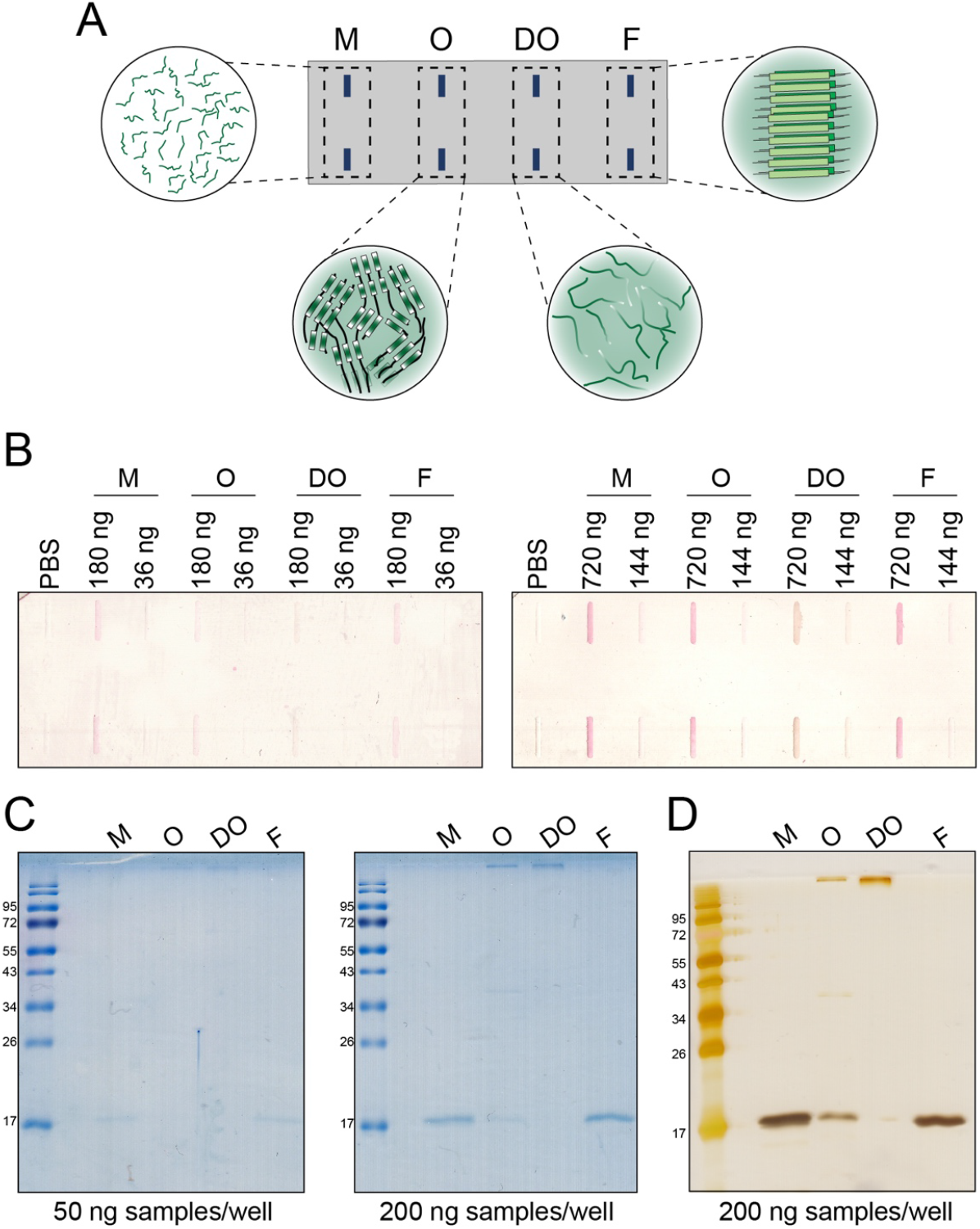
A) A Schematic illustration of slot blot performed with different α-syn monomers (**M**, unstructured), β-sheet rich oligomers (**O**), dopamine-induced unstructured oligomers (**DO**) and fibrils (**F**, β-sheet rich). B) Ponceau S staining on nitrocellulose membranes loaded with different α-syn samples used at varying concentration to ensure equal loading on the membranes for slot blot experiments. C and D) SDS-PAGE analysis followed by Coomassie staining (C) and silver staining (D) of α-syn samples (monomers, O, DO and fibrils) Abbreviations: M: Monomers; O: Unmodified Oligomers; DO: Dopamine-induced oligomers; F: Fibrils.

**Supplementary figure 2:**
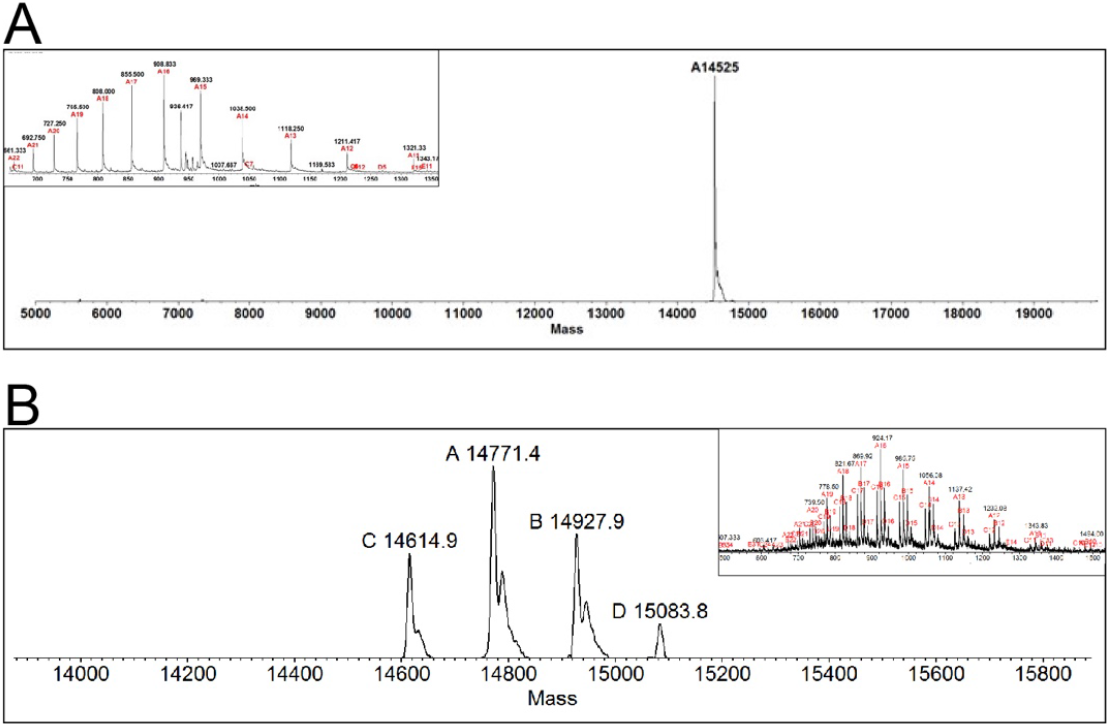
Mass spectrometry analysis of α-syn monomers purified by size exclusion chromatography. α-syn monomers purified from preparation of Dopamine-induced oligomers(A) and HNE-induced oligomers (B).

**Supplementary figure 3:**
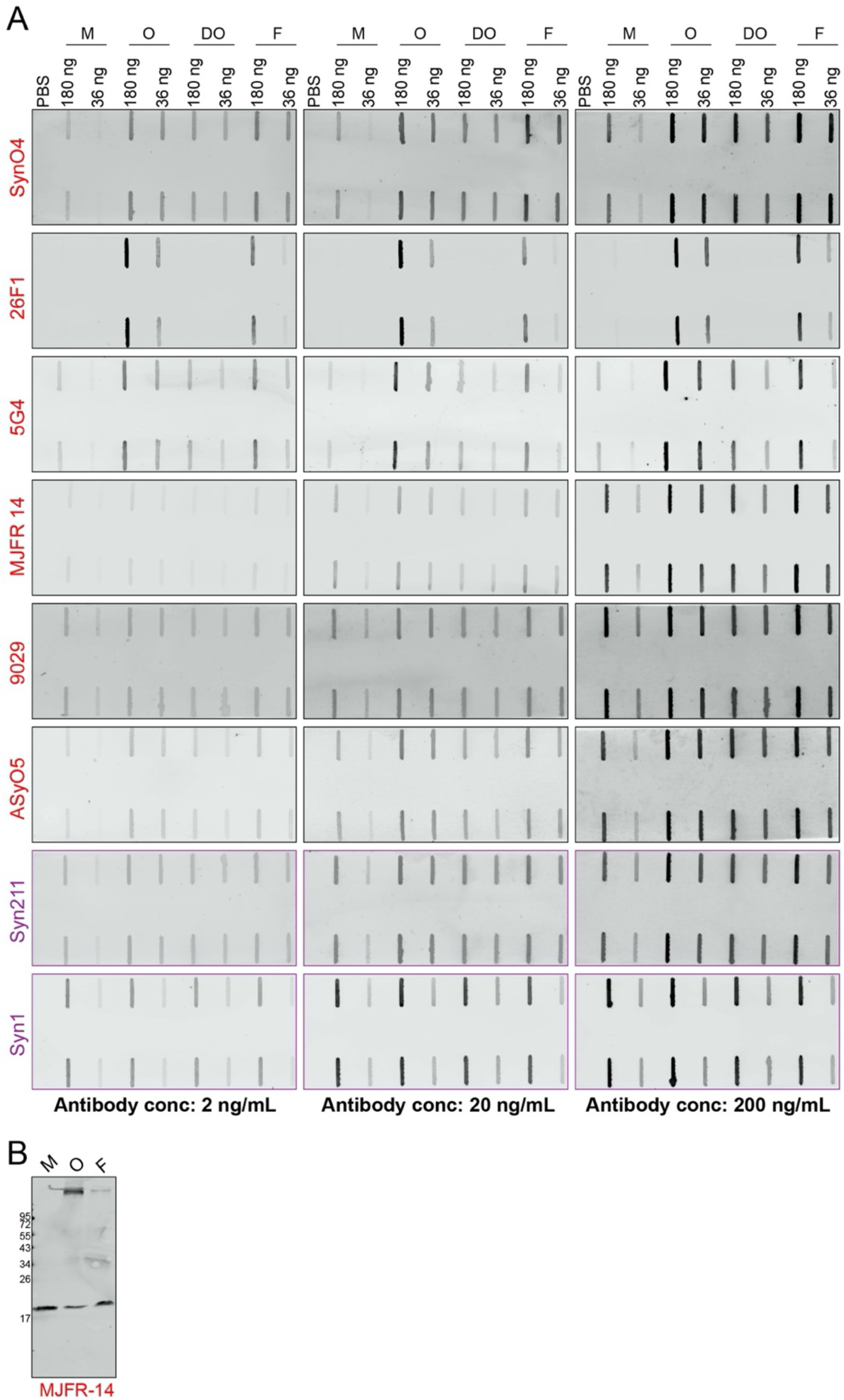
(A) Specificity of antibodies against α-syn monomers (M), unmodified oligomers (O), Dopamine-induced oligomers (DO), and fibrils (F) using increasing concentrations of antibodies (2 ng/mL, 20 ng/mL, 200 ng/mL and). **(B) WB** assessment of the immunoreactivity of MJFR-14 (2 ng/mL) against α-syn samples (180 ng).

**Supplementary figure 4:**
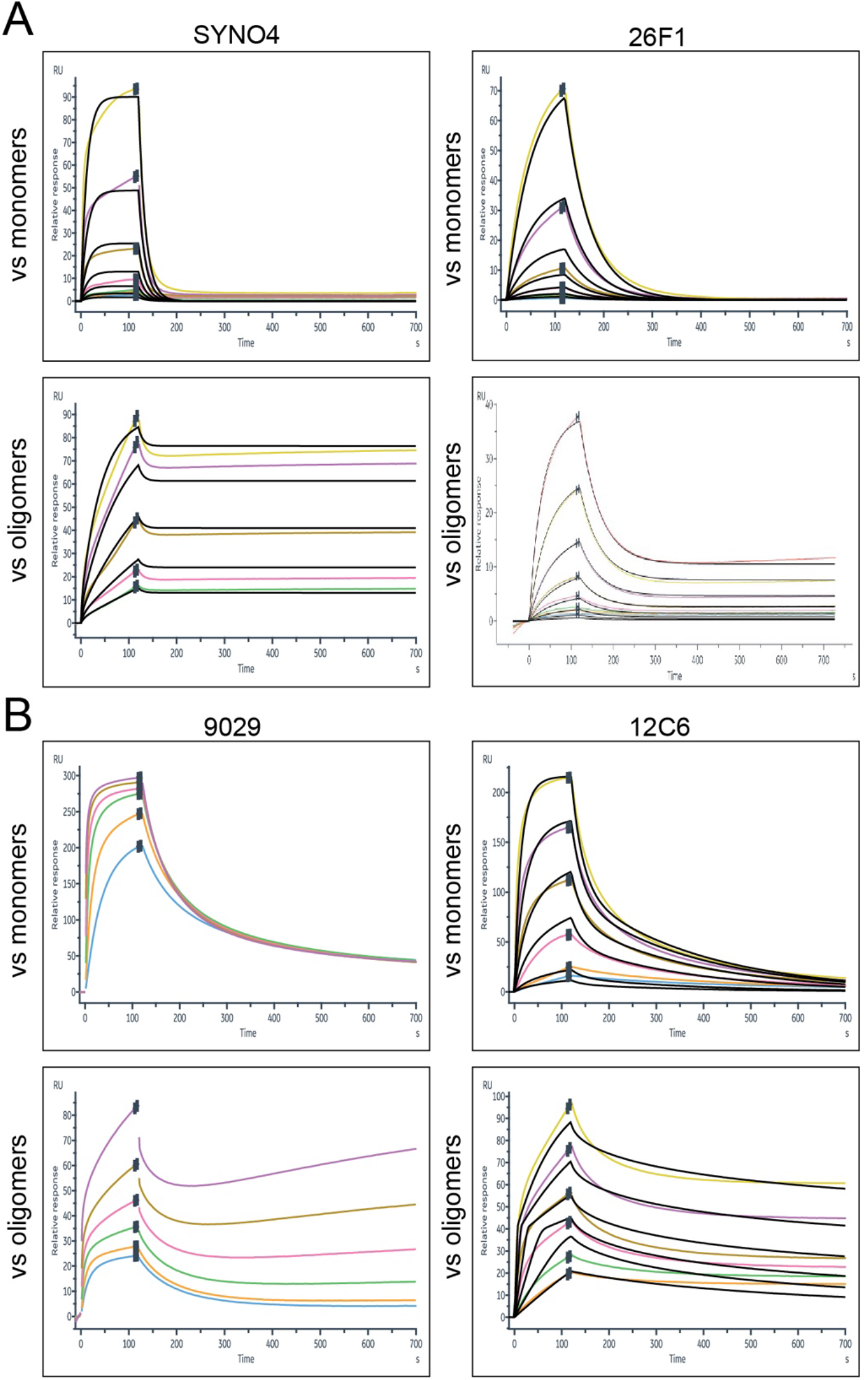
SPR-based kinetic analysis of different immobilized monoclonal antibodies (SynO4, 26F1, 9029 and 12C6) binding to α-syn monomers (A) and unmodified oligomers (B).

